# Human Early Syncytiotrophoblasts Are Highly Susceptible to SARS-CoV-2 Infection

**DOI:** 10.1101/2022.11.17.516978

**Authors:** Degong Ruan, Zi-Wei Ye, Shuofeng Yuan, Zhuoxuan Li, Weiyu Zhang, Chon Phin Ong, Kaiming Tang, Jilong Guo, Yiyi Xuan, Timothy Theodore Ka Ki Tam, Yunying Huang, Qingqing Zhang, Cheuk-Lun Lee, Philip C.N. Chiu, Fang Liu, Dong-Yan Jin, Pentao Liu

## Abstract

The ongoing and devastating pandemic of coronavirus disease 2019 (COVID-19) has led to a global public health crisis. COVID-19 is caused by severe acute respiratory syndrome coronavirus 2 (SARS-CoV-2) and can potentially pose a serious risk to maternal and neonatal health. Cases of abnormal pregnancy and vertical transmission of SARS-CoV-2 from mother to foetus have been reported but no firm conclusions are drawn. Trophoblasts are the major constituents of the placenta to protect and nourish the developing foetus. However, direct *in vivo* investigation of trophoblast’s susceptibility to SARS-CoV-2 and of COVID-19 and pregnancy is challenging. Here we report that human early syncytiotrophoblasts (eSTBs) are highly susceptible to SARS-CoV-2 infection in an angiotensin-converting enzyme 2 (ACE2)-dependent manner. From human expanded potential stem cells (hEPSCs), we derived *bona fide* trophoblast stem cells (TSCs) that resembled those originated from the blastocyst and the placenta in generating functional syncytiotrophoblasts (STBs) and extravillus trophoblasts (EVTs) and in low expression of HLA-A/B and amniotic epithelial (AME) cell signature. The EPSC-TSCs and their derivative trophoblasts including trophoblast organoids could be infected by SARS-CoV-2. Remarkably, eSTBs expressed high levels of ACE2 and produced substantially higher amounts of virion than Vero E6 cells which are widely used in SARS-CoV-2 research and vaccine production. These findings provide experimental evidence for the clinical observations that opportunistic SARS-CoV-2 infection during pregnancy can occur. At low concentrations, two well characterized antivirals, remdesivir and GC376, effectively eliminated infection of eSTBs by SARS-CoV-2 and middle east respiratory syndrome-related coronavirus (MERS-CoV), and rescued their developmental arrest caused by the virus infection. Several human cell lines have been used in coronavirus research. However, they suffer from genetic and/or innate immune defects and have some of the long-standing technical challenges such as cell transfection and genetic manipulation. In contrast, hEPSCs are normal human stem cells that are robust in culture, genetically stable and permit efficient gene-editing. They can produce and supply large amounts of physiologically relevant normal and genome-edited human cells such as eSTBs for isolation, propagation and production of coronaviruses for basic research, antivirus drug tests and safety evaluation.

## INTRODUCTION

The possibility of SARS-CoV-2 infection during pregnancy has direct implications for the mother and foetus. The human placenta protects the fetus using multiple cellular and molecular defense mechanisms at the maternal-foetal interface safeguarding against SARS-CoV-2 infection during pregnancy. Notwithstanding, SARS-CoV-2 infection of the human placenta and resultant damages have been reported that caused placental inflammation, trophoblast necrosis and chronic histiocytic intervillositis (Algarroba et al., 2020; Hosier et al., 2020; Schwartz and Morotti, 2020; Vivanti et al., 2020). In rare cases of pregnant COVID-19 patients, intrauterine vertical transmission from mother to foetus was found or suspected (Baud et al., 2020; Buonsenso et al., 2020; Dong et al., 2020b; Penfield et al., 2020; Vivanti et al., 2020; Zamaniyan et al., 2020; Zeng et al., 2020) although these cases were often in the late stages of pregnancy with the possible confounding of postpartum infection (Kreis et al., 2020; Raschetti et al., 2020).

The human placenta consists of both maternal and foetal tissues (Burton and Fowden, 2015). The extraembryonic ectoderm in early post-implantation embryos generates the proliferative mononucleated trophoblast progenitors (villous cytotrophoblasts (vCTBs), which can differentiate into invasive EVTs in the anchoring villi that grow out into the maternal decidua. vCTBs can also undergo active cellular fusion into non-proliferative multinucleated STBs that form the layer covering placenta villi. This STB layer presents a physical barrier to pathogens since it lacks intercellular gap junctions that can be exploited by pathogens or modulated by inflammatory signals (Zeldovich et al., 2013).

SARS-CoV-2 infects cells via its spike (S) protein binding the host entry receptor ACE2 (Letko et al., 2020; Zhou et al., 2020) primed by the human transmembrane protease serine 2 (TMPRSS2) (Hoffmann et al., 2020). Molecular assays and single-cell RNA sequencing (scRNAseq) studies have identified ACE2 and TMPRSS2 expression in human placenta cells at several gestation stages, including vCTBs, STB and EVTs (Ashary et al., 2020; Li et al., 2020; Liu et al., 2018; Vento-Tormo et al., 2018; Zheng et al., 2020), indicating the potential susceptibility of trophoblasts. However, term placenta trophoblasts are not susceptible to SARS-CoV-2 and do not express ACE2 (Colson et al., 2021), and it is technically challenging to obtain normal trophoblasts of earlier pregnancy stages to test virus infection. Furthermore, laboratory model organisms such as the mouse have substantial differences from humans in trophoblast biology and placenta development, therefore 2D and 3D cellular models of normal human trophoblasts are needed to decipher SARS-CoV-2 infection in trophoblasts and in pregnancy.

Expanded potential stem cells (EPSCs) are a new stem cell type derived from cleavage stage preimplantation embryos and retain developmental potential to both extra-embryonic as well as embryonic cell lineages (Gao et al., 2019; Yang et al., 2017a; Yang et al., 2017b). In particular, hEPSCs directly generated hTSCs *in vitro* (Cinkornpumin et al., 2020; Gao et al., 2019). Although standard human embryonic stem cells (hESCs) could be induced to generate trophoblast-like cells (Amita et al., 2013; Horii et al., 2016; Xu et al., 2002), these cells however did not fulfil stringent criteria for trophoblast identity (Bernardo et al., 2011; Lee et al., 2016b; Roberts et al., 2018). Human naïve ESCs were recently shown to directly give rise to hTSCs, as confirmed by morphological, molecular and transcriptomic criteria (Cinkornpumin et al., 2020; Dong et al., 2020a; Guo et al., 2021), which may reflects the unique property of human naïve epiblast to regenerate trophoblast (Guo et al., 2021). However, naïve ESCs are known to have genome-wide DNA demethylation and potential genomic imprinting defects and genetic instability (Choi et al., 2017; Theunissen et al., 2014; Yagi et al., 2017). Some human naïve cell cultures may already have trophoblasts (Messmer et al., 2019) although an improved naïve cell condition could reduce point mutation load (Stirparo et al., 2021).

In the present study, we took advantage of hEPSC’s potent capability to generate trophoblasts and established and validated a stem cell-based system to interrogate trophoblast’s SARS-CoV-2 susceptibility. SARS-CoV-2 infected EPSC-derived hTSCs, EVTs, and STBs but with relatively low efficiencies. In stark contrast, early STBs or eSTBs generated from hTSCs expressed high levels of ACE2 and were highly potent in SARS-CoV-2 infection and virus production. Genetic knockout of ACE2 abolished SARS-CoV-2 infection of trophoblasts. Low concentrations (nM) of FDA-approved antiviral drugs, remdesivir and GC376, also effectively eliminated eSTB viral infection. hEPSCs have a broader developmental potential, are robust in culture and genetically stable, and enables efficient gene-editing (Gao et al., 2019; Yang et al., 2017a; Zhao et al., 2021). These stem cells therefore may serve to identify susceptible cell types and to study host factors in coronavirus infection. They can provide new cell sources such as eSTBs for coronavirus isolation, propagation, and production, and help address some of the challenging technical issues of the currently used mammalian cells in virus research.

## RESULTS

### Establishment of EPSC-TSCs and generation of STBs and EVTs for infection

In order to develop hEPSCs as a model for COVID-19 study, we established several new human trophoblast stem cell (hTSC) lines using the published condition (Okae et al., 2018) from M1 hEPSCs, which were converted from primed human ESC line M1 (Gao et al., 2019) (Figures 1A and 1B). These new EPSC-TSCs would also enable further confirmation the identity of hTSCs since it was recently found that hEPSCs also generated AME cells together with trophoblasts (Gao et al., 2019; Guo et al., 2021)..

**Figure 1.**
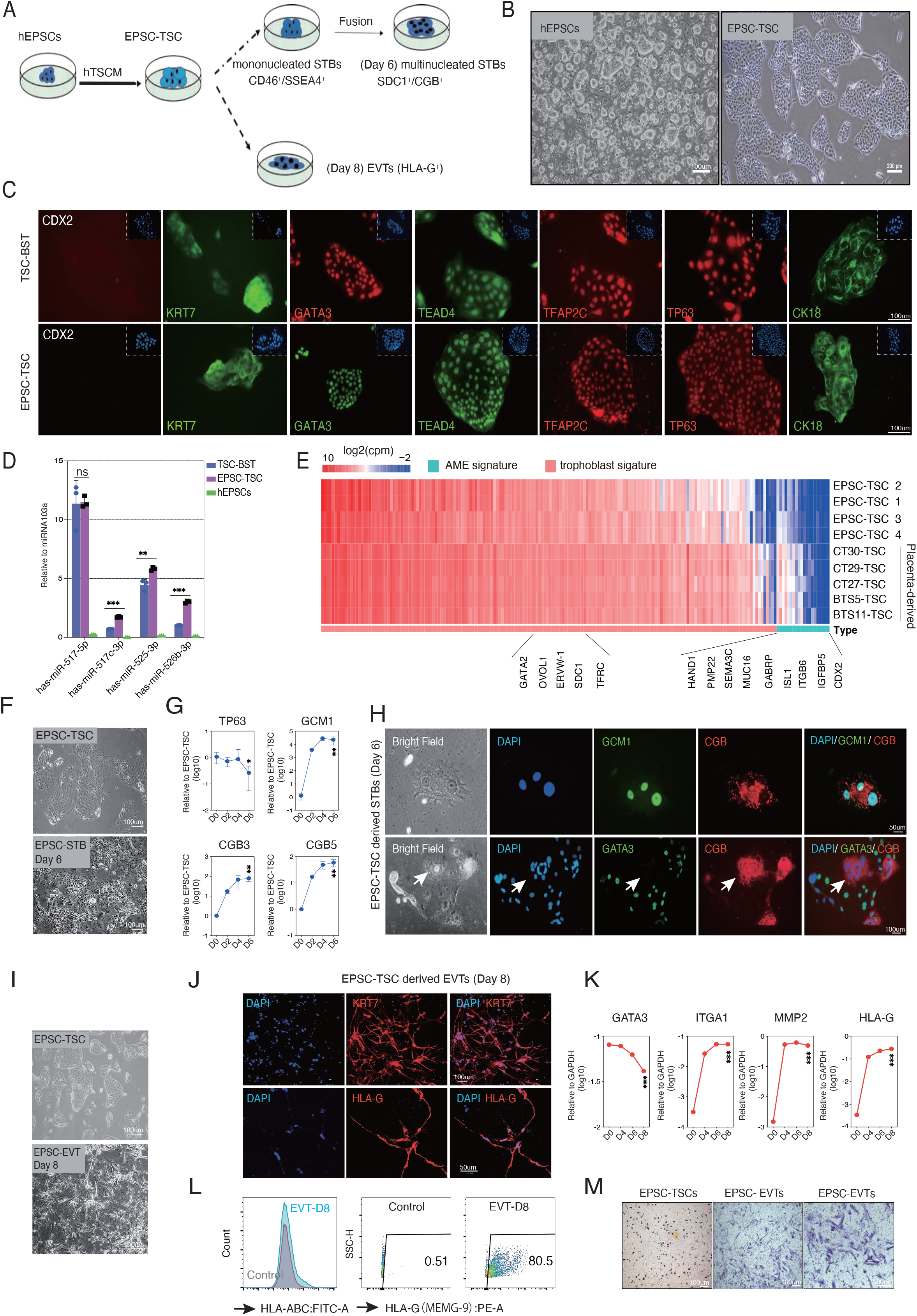
Generating trophoblast stem cells (TSCs) and subtype trophoblasts from hEPSCs. (A) Experiment flow of sequential generation of hTSCs, STBs, and EVTs from hEPSCs. Under the hTSC medium (hTSCM), EPSC-TSCs are directly established from hEPSCs. EPSC-TSCs are induced to differentiate into STBs and EVTs. The initially differentiated EPSC-TSCs toward STBs are the mononucleated cells that are positive for early STB markers SSEA4 and CD46, while the late-stage differentiated cells are multinucleated and express more mature STB markers SDC1 and CGB. EPSC-TSCs are also induced to generate EVTs that express HLA-G on day 8 of differentiation. (B) Images of hEPSCs and EPSC-TSCs. Scale bar: left panel, 100μm; right panel, 200μm (C) EPSC-TSCs and human blastocyst derived hTSCs (TSC-BST) (Okae et al., 2018) are immunofluorescence stained for CDX2 and hTSC markers KRT7, GATA3, TEAD4, TFAP2C, TP63, CK18. Both EPSC-TSC and TSC-BST are negative for CDX2. DAPI stains the nucleus. Scale bar: 100μm (D) RT-qPCR detection of the trophoblast-specific C19MC miRNAs (has-miR-517a, 517b, 525-3p and 526-3p) in hEPSCs, EPSC-TSCs and TSC-BST. Results are normalized to levels of miR-103a using the ΔCt method. Three independent experiments were performed. Data are mean ± SD. ns, not significant; **p<0.01; *** p < 0.001 (two-tailed unpaired Student’s t-test). (E) RNAseq analysis of trophoblast and AME genes (Table S1) expressed in EPSC-TSCs and in five human placenta-derived hTSCs (CT27, 29, 30 and BTS5 and 11) (Sheridan, M.A., 2021). Log2 transformed cpm (counts per million) are used in generating the heatmap. Individual genes of the two gene sets are ordered by expression levels. All hTSCs express high levels of trophoblast genes and low levels of amniotic lineage genes. (F) EPSC-TSC differentiation toward STBs on day 6 (STB-D6). Scale bar: 100μm (G) RT-qPCR detects decreased expression of TP63 and increased STB markers CGB5, GCM1 and CGB3 in EPSC-TSC differentiation toward STBs. STB-D2, -D4 and -D6 are days along the differentiation. Results are normalized to levels of GAPDH using the ΔCt method. Three independent experiments were performed. Data are mean ± SD. *p<0.05; ** p < 0.01 (one-tailed unpaired Student’s t-test between EPSC-TSCs and STB-D6). (H) EPSC-TSC differentiation toward STBs on day 6 and immunofluorescence staining images for STB markers GCM1 and CGB (upper panel); and GATA3 (lower panel). Arrows indicate the lack of GATA3 in a multinucleated CGB^+^ mature STB. DAPI stains the nucleus. Scale bar: top panel 50 μm; lower panel 100 μm (I) EVTs generated on day 8 of EPSC-TSC differentiation. Scale bar: 100μm (J) EVTs generated from EPSC-TSCs express KRT7 and EVT specific marker HLA-G in immunofluorescence staining. DAPI stains the nucleus. Scale bar: top panel 100 μm; lower panel 50μm (K) RT qPCR detects decreased expression of GATA3 and increased EVT markers ITGA1, MMP2 and HLA-G in EPSC-TSC differentiation toward EVTs. EVT-D2, -D4 and -D6 are days along the differentiation. Results are normalized to levels of GAPDH using the ΔCt method. Three independent experiments were performed. Data are mean ± SD. *** p < 0.001 (two-tailed unpaired Student’s t-test between EPSC-TSCs and EVT-D8). (L) Flow cytometry quantification of expression of HLA-A,B,C and EVT specific marker HLA-G on EVTs generated from EPSC-TSCs. EPSC-TSCs were used as the flow analysis control. Both TSCs and EVTs are negative for the HLA-A,B,C antibody. (M) Evaluation of invasiveness of EVTs generated from EPSC-TSCs in transwell assay. The control EPSC-TSCs have little invasive activity. Scale bar: 100μm

EPSC-TSCs expressed typical hTSC factors such as TFAP2C, TP63, CK18, GATA3, ELF5, TEAD4 and KRT7 (Figure 1C and Figure S1A), and trophoblast-specific miRNAs C19MC miRNAs (has-miR-517c-3p, 517-5p, 525-3p and 526b-3p) (Lee et al., 2016a) (Figure 1D), highly similar to human blastocyst-derived TSCs (TSC-BST) (Okae et al., 2018). They were low or negative for the classical HLA class I molecules HLA-A and -B, also like TSC-BST (Figures S1B and S1C). In both RNAseq analysis and individual gene RT-PCR, EPSC-TSCs did not express those currently recognized AME signature genes such as CDX2, MUC16, GABRP, ITGB6, or VTCN1 (Zheng et al., 2019) (Figure 1E, Figures S1D and S1E, and Table S1), resembling the recently published hTSCs derived from human placenta trophoblasts (CT27, 29, 30-TSC) (Sheridan et al., 2021).

EPSC-TSCs were induced to generate STBs (Figures 1F and S1F). Trophoblast progenitor transcription factor TP63 was reduced whereas STB genes such as GCM1, β chorionic gonadotrophin 3 (CGB3) and CGB5 was quickly increased (Figure 1G). Immunofluorescence staining of day 6 differentiated cells (STB-D6) detected GCM1^+^ and CGB^+^ and multinucleated STBs, which appeared to have low GATA3 levels (Figure 1H and S1G). Functionally, in the supernatant of STB-D6 cells, ELISA detected properly folded and secreted β-hCG hormone (Figure S1H). Under TGFβ inhibition, EPSC-TSCs could efficiently generate EVTs, which had the typical EVT morphology and were stained positively for KRT7, HLA-G, ITGA1 and IGTA5 (Figures 1I-1J, Figure S1I). EVT genes ITGA1, MMP2 and HLA-G were rapidly increased whereas GATA3 was reduced (Figure 1K). Indeed, GATA3 was not detectable in HLA-G^+^ EVTs (Figure S1J). On day 8, most cells were positive for HLA-G (80%), ITGA1 (82%) and ITGA5 (95%), but negative for the classical HLA class I molecules, HLA-A and -B (Figure 1L and Figure S1K). Functionally, EPSC-TSC derived EVTs possessed potent invasiveness capability (Figure 1M).

We next performed RNAseq analysis of EPSC-TSCs and their trophoblast derivatives STBs and EVTs, which expressed distinct sets of genes specific for human primary trophoblast progenitor, STB and EVT (Sheridan et al., 2021) (Figure S1L). RNAseq also confirmed low classical HLA class I molecules HLA-A and -B (S1M) as in those originated from the placenta trophoblasts (Sheridan et al., 2021).

### Trophoblasts derived from hEPSCs molecularly resemble those in the human peri-implantation embryos and the placenta

To further validate the identity of the *in vitro* generated trophoblasts, we directly compared EPSC-TSCs and their derivatives to trophoblasts in peri-implantation embryos and in the placenta. We first extracted scRNA-seq data of 4041 peri-implantation extra-embryonic cells (embryonic day 6 to 14) in the prolonged culture of human embryos (Zhou et al., 2019) and 952 placental cells from first and second trimester pregnancies (Liu et al., 2018), and subsequently combined these scRNA-seq datasets and computed the joint uniform manifold approximation and projection (UMAP), with their developmental times highlighted (in embryonic day or ED, or gestational week) (Figure 2A) and developmental stage and subtype annotated (Figure 2B). The peri-implantation sector contains the trophoblast cells (PI-TB) which possess stemness and the potency to differentiate into EVT (PI-EVT) and STB (PI-STB) as shown in the same sector (Figure 2B). Correspondingly, the placenta cells can be identified as either first-trimester cytotrophoblast cells (T1-CTB), T1-EVT and T1-STB, or second-trimester placenta EVT (T2-EVT) (Figure 2B). The trophoblast marker genes were examined in each stage-specific subtypes, which validated their trophoblast identities (Figure S2A). Notably, these human *in vivo* trophoblasts did not appear to highly express the reported amnion markers MUC16, GABRP or CDX2, whereas some did express VTCN1, ITGB6 and ISL1 (Figure S2B).

**Figure 2.**
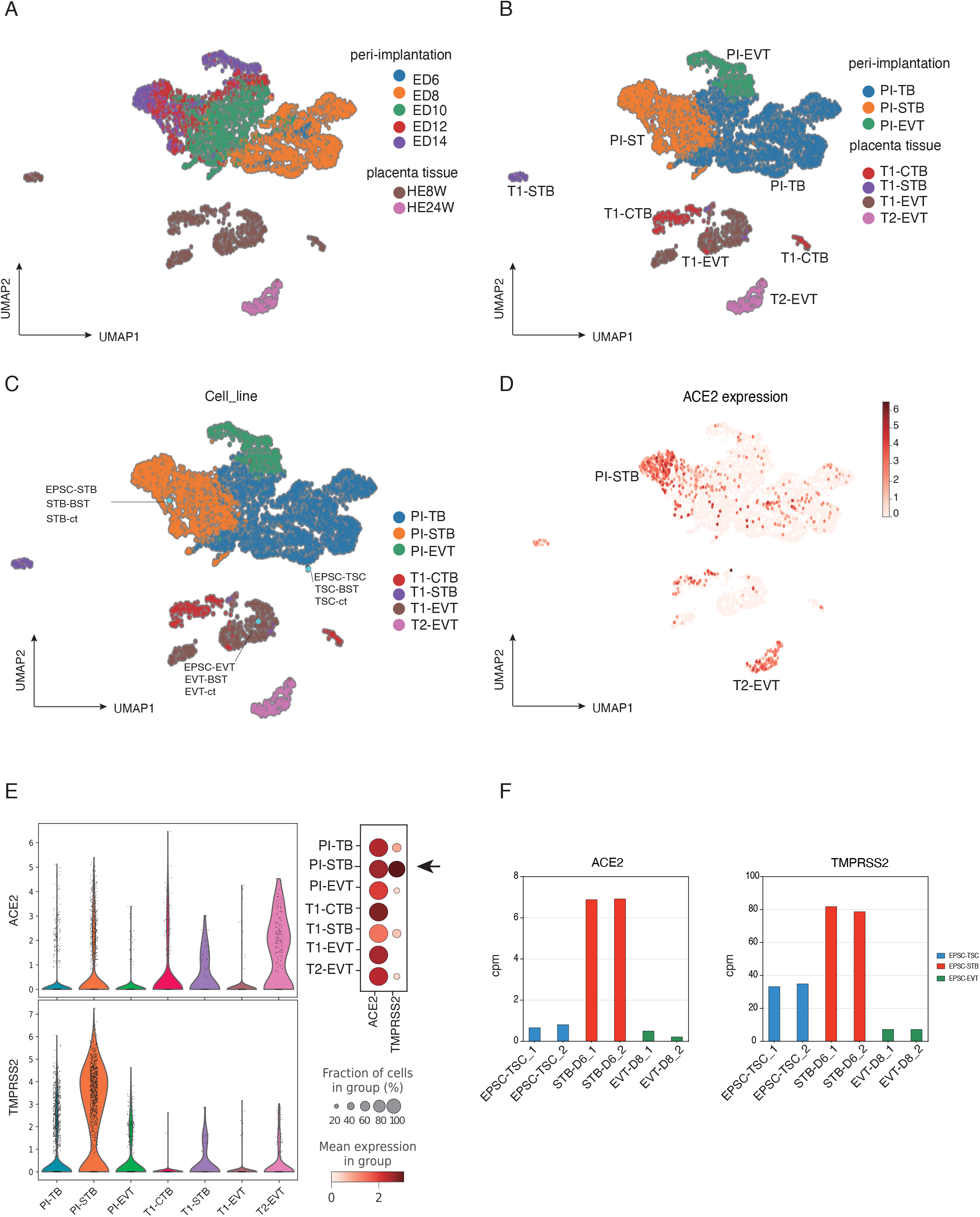
Trophoblasts derived from hEPSCs resemble those in the human embryos and the placenta. (A-B) Umap representations containing scRNAseq data of cells from *in vitro* cultured peri-implantation embryo stages (top panel) and of cells from first and second trimester placenta (bottom panel). Cells are colored by developmental time points (ED6-14: Embryonic day 6-14. HE8W / HE24W: placenta trophoblasts at 8 weeks or 24 weeks of gestation, corresponding to first or second trimester) in (A), and trophoblast types at each stage in (B). PI: Peri-implantation; PI-TB: Peri-implantation trophoblast; PI-STB: Peri-implantation syncytiotrophoblast; PI-EVT: Peri-implantation extravillus trophoblast; T1-CTB, T1-STB and T1-EVT: first trimester cytotrophoblast, syncytiotrophoblasts and extravillus trophoblast; T2-EVT: second trimester extravillus trophoblast. (C) Mapping *in vitro* trophoblast cells (EPSC-TSCs, TSC-BST and TSC-ct, and their derivatives) bulk RNA-seq data to the peri-implantation and placenta trophoblast clusters in Figure 2B. *In vitro* cells are highlighted in light blue. TSC-BST and TSC-ct are derived from the human blastocyst and placenta cytotrophoblasts as reported (Okae et al., 2018). STB-BST, STB-ct, and EPSC-STB, and EVT-BST, EVT-ct, and EPSC-EVT are derived from the respective TSCs. (D) Expression of ACE2 in peri-implantation and placenta trophoblast scRNAseq clusters. ACE2 is expressed in many PI-STBs. (E) Left panel: Violin plots show ACE2 and TMPRSS2 expression levels (log-transformed TPM) detected in scRNAseq dataset among different lineages at different stages in human peri-implantation embryo and placenta tissue. Right: Dot plot for ACE2 and TMPRSS2 expression levels in ACE2-positive cells, categorized according to stage and cell lineage. Dots are colored according to the mean expression value in each category and dot size indicates the percentage of ACE2- or TMPRSS2-positive cells from each category that expresses ACE2. Only PI-STBs express high levels of both ACE2 and TMPRSS2. (F) RNA-seq expression of ACE2 and TMPRSS2 in EPSC-TSCs and the differentiated subtype cells. STB-D6 and EVT-D8 are differentiated cells from EPSC-TSCs on day 6 and day 8, respectively. Cpm was used to normalize the signal.

We then projected and mapped EPSC-TSCs and STB and EVT derived from them, and blastocyst- or cytotrophoblast-derived TSCs (TSC-BST/ct) and their STB/EVT (Okae et al., 2018), against the *in vivo* trophoblast clusters categorized by stage and lineage as described in Figure 2A-2B. All hTSCs (EPSC-TSC, TSC-BST, TSC-ct) were projected in a proximity to PI-TB, which possess stemness, whereas STB cells to PI-STB and EVT cells to T1-EVT, respectively, regardless of cell line origins (Figure 2C). The resemblance between *in vitro* generated trophoblasts and the *in vivo* ones was supported by whole-transcriptome Pearson correlation (Figure S2C).

To explore potential trophoblast susceptibility to SARS-CoV-2, we first examined SARS-CoV-2 host factor genes in the *in vivo* trophoblasts. The prevalently recognized SARS-CoV-2 receptor ACE2 expression was high in many cells in the PI-STB cluster (Figures 2D and 2E). Correlation analysis between ACE2 and trophoblast subtype markers only detected significant positive correlations with STB markers such as CD46, CGB5, ENG, and CSH2 (r = 0.245, 0.200, 0.207, 0.248, respectively, p < 0.0001) (Figure S2D) but with not EVT genes (Table S2). In the *in vivo* trophoblast datasets, only the PI-STB cluster expressed high levels of both ACE2 and TMPRSS2 (Figure 2E and Figure S2E). In contrast to ACE2 and TMPRSS2, other reported SARS-CoV-2 receptor-related genes such as BSG and AXL, did not appear to be very specifically expressed in a particular cell cluster (Figure S2E).

In the *in vitro* cultured trophoblasts, EPSC-TSCs, TSC-BST/ct, and the STBs and EVTs derived from them, also expressed ACE2 and TMPRSS2 with STBs having the highest levels (Figure 2F, and Figures S2F-S2G). The host factor gene expression profiles in both *in vivo* and *in vitro* human trophoblasts implicate their susceptibility to SARS-CoV-2 infection.

### eSTBs are highly susceptible to SARS-CoV-2 infection

To experimentally investigate trophoblast susceptibility to SARS-CoV-2, we subjected hEPSCs, EPSC-TSCs and the derived STBs and EVTs to SARS-CoV-2 infection. Briefly, cells were co-cultured with SARS-CoV-2 (SARS-CoV-2 HKU-001a strain; GenBank accession number MT230940) for 2 hrs followed by incubation in fresh medium for another 24, 48, or 72 hrs (Figure 3A). The supernatant and cell lysates were collected for viral genome, protein and live virus detection. hEPSCs did not express ACE2 (Figure S2G) thus were not infectable by SARS-CoV-2. No substantially increased detection of the viral genome in the supernatant or cell lysate, and no viral N protein was visible after immunofluorescence staining (Figure 3B, and Figures S3A and S3B). In line with relatively low ACE2 and TMPRSS2 levels, SARS-CoV-2 did not efficiently infect EPSC-TSCs or EVTs, with only about 3-4% or 2% cells being positive for the viral N protein (Figures 3B-3D, and Figures S3A, S3C-S3E), respectively.

**Figure 3.**
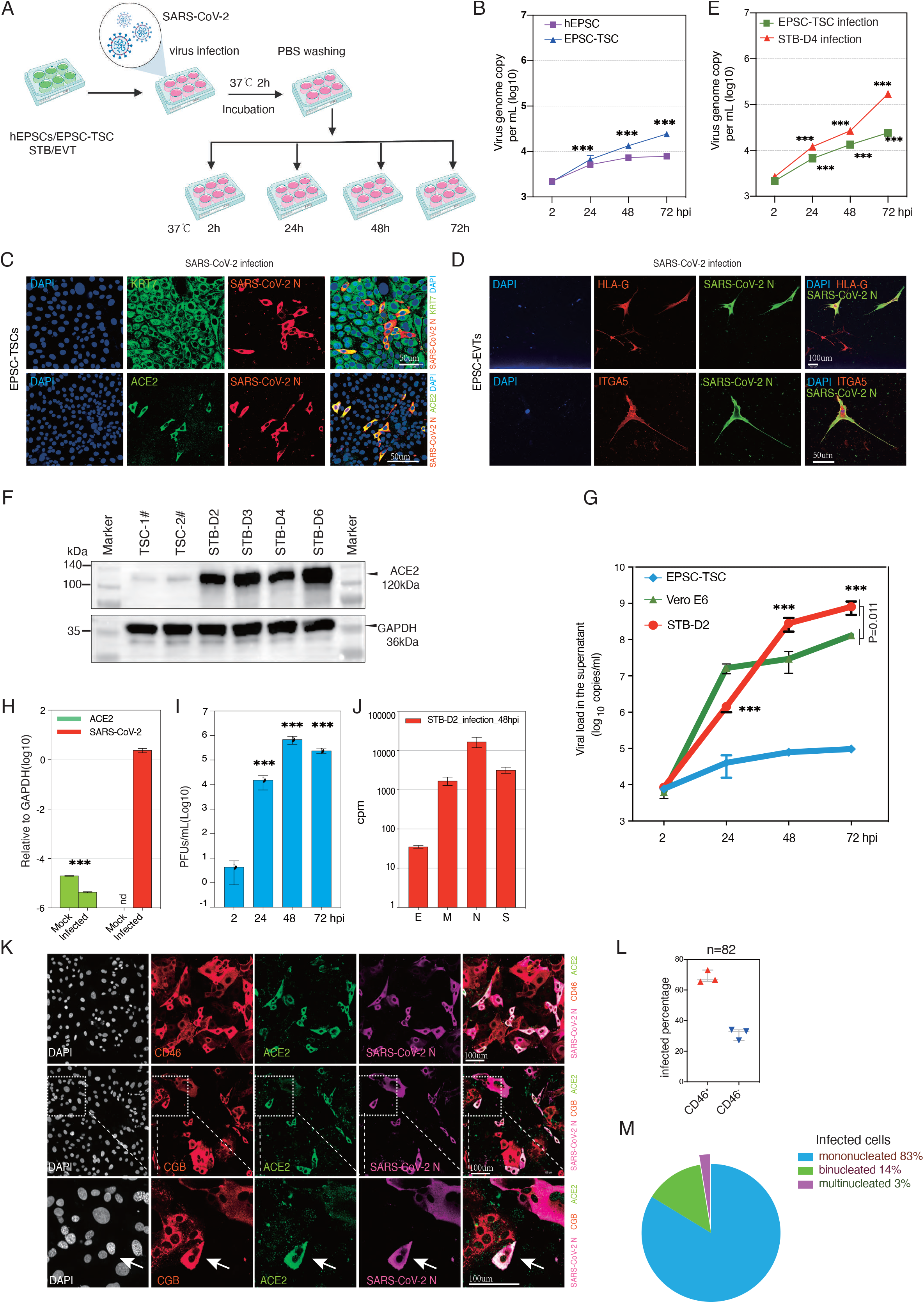
eSTBs are highly susceptible to SARS-CoV-2 infection. (A) Experimental flow of SARS-CoV-2 infection of trophoblasts. Cells were co-cultured with SARS-CoV-2 at 0.1 MOI for 2 hrs followed by PBS wash, and were subsequently incubated in fresh trophoblast media for 24, 48, or 72 hrs. The supernatants and cell lysates were collected for viral genome detection and viral protein analysis. All the virus infection experiments were repeated at least three times. (B) RT-qPCR detection of SARS-CoV-2 genome copy numbers (per mL) in the supernatants of virus-co-cultured hEPSCs and EPSC-TSCs at different time points. Datapoints are mean and SD from three independent experiments. *** p < 0.001 (two-tailed unpaired Student’s t-test). (C) Detection of KRT7, ACE2 and SARS-CoV-2 N protein in SARS-CoV-2 infected EPSC-TSCs. Only ACE2-expressing cells are infected. Cells were fixed for staining immunofluorescence staining 48 hours after virus co-incubation. DAPI stains the nucleus. Scale bar: 50μm (D) SARS-CoV-2 infected EVTs stained for N protein and EVT markers HLA-G and ITGA5 in immunofluorescence staining. DAPI stains the nucleus. Scale bar: top panel 100 μm; lower panel 50μm (E) RT-qPCR detection of SARS-CoV-2 genome copy numbers (per mL) in the supernatants of virus-co-cultured EPSC-TSCs and STBs. The STBs were differentiated from EPSC-TSCs for 4 days (STB-D4) before SARS-CoV-2 infection. Datapoints are mean and SD from three independent experiments. *** p < 0.001 (two-tailed unpaired Student’s t-test). (F) Detection of ACE2 protein in EPSC-TSCs and during their differentiation toward STBs. ACE2 expression is rapidly induced and at high levels in STB-D2 (day 2) and onward. (G) RT-qPCR detection of SARS-CoV-2 genome copy numbers (per mL) in the supernatants of virus-co-cultured EPSC-TSCs and eSTBs (STB-D2). Supernatants were harvested at several time points post co-incubation. Vero E6 cells are widely used in SAS-CoV-2 research and were used as a control in virus genome amplification. Datapoints are mean and SD from three independent experiments. *** p < 0.001 (two-tailed unpaired Student’s t-test). P=0.011 (one-tailed unpaired Student’s t-test between STB-D2 72 hpi and Vero E6 72 hpi). (H) RT-qPCR detection of ACE2 and SARS-CoV-2 N gene expression in cell lysates of the infected eSTBs (STB-D2) at 48 hpi. nd, not detectable. Data are mean ± SD. *** p < 0.001 (two-tailed unpaired Student’s t-test). (I) Plaque formation assay to detect SARS-CoV-2 virus in the supernatants from SARS-CoV-2-inoculated STB-D2 (48 hpi). Data were obtained from three (n=3) independent batches of STB-D2. Datapoints are mean and SD from three independent experiments. *** p < 0.001 (two-tailed unpaired Student’s t-test). PFUs: plaque forming units (J) RNAseq analysis of SARS-CoV-2 genes (E, M, N, S) in cell lysates of 48 hpi eSTBs (STB-D2). (K) Representative immunofluorescence staining images of 24 hpi eSTBs (STB-D2) for ACE2 and SARS-CoV-2 N protein, and for STB markers CD46 and CGB. The infected cells are positive for ACE2 and CD46, but rarely multinucleated CGB^+^ cells. Arrows indicate an infected binucleated STB. DAPI stains the nucleus. Scale bar: 100μm. (L) Quantification of the percentages of early STB marker CD46 in 24 hpi eSTBs (STB-D2). Values are mean ± SEM. (M) Quantification of the percentages of mono-, bi-, and multi-nucleated cells in 24 hpi eSTBs (STB-D2). SARS-CoV-2 preferably infects mononucleated cells.

In contrast to EPSCs, EPSC-TSCs and EVTs, STBs expressed high levels of both ACE2 and TMPRSS2, and were more efficiently infected by SARS-CoV-2 (Figure 3E and Figure S3F). We noticed that the virus-infected cells were often positive for eSTB markers (SSEA4 and CD46) (Gamage et al., 2016; Holmes et al., 1992) but didn’t appear to highly express mature STB gene CGB (Figure S3G). Indeed, multinucleated and CGB^+^ mature STBs did not appear to express high levels of ACE2 (Figure S3G) and accounted for only a minor population of the infected cells (Figure S3H). These experimental results indicate that SARS-CoV-2 more efficiently infects immature or early STBs compared to multinucleated mature STBs.

To directly investigate STBs susceptibility for SARS-CoV-2, we examined STBs at several time points following induction of EPSC-TSC differentiation toward STB. In Day 2 differentiation (STB-D2), most cells were mononucleated and negative for CGB. Only 3% of cells were multinucleated (Figure S3I-S3J). The eSTB gene CD46 was transiently up-regulated in STB-D2 (Figure S3K). Notably, ACE2 expression was rapidly upregulated in STB-D2, together with TMPRSS2 (Figure 3F and Figures S3L-S3M). We thus used STB-D2 cells for the subsequent infection experiments and empirically named these cells as eSTBs.

Following STB-D2 or eSTB cell infection with SARS-CoV-2, supernatants were collected at 2, 24, 48 and 72 hrs and quantified for viral RNA loads. Compared to Vero E6 cell infection, eSTBs produced around 6 folds more supernatant viral RNA at 48- and 72-hour post-infection (Figure 3G). Substantial amounts of viral genomes were also detected in cell lyses (Figure 3H). Plaque assay confirmed a substantial amount of infectious virus particles released from the infected eSTBs (Figure 3I). The potent production of SARS-CoV-2 was further revealed in RNAseq where abundant transcripts of viral Envelope (E), Membrane glycoprotein (M), Nucleocapsid (N) and Spike (S) genes were detected (Figure 3J). Immunostaining for SARS-CoV-2 N protein revealed that the virus infected cells were primarily CD46^+^ whereas those multinucleated cells were rarely infected (Figures 3K-3M). The efficient virus propagation and production in eSTBs, which are derived from normal human stem cells, demonstrate that eSTBs can serve as host cells for studying infectious disease.

### SARS-CoV-2 infection impairs EPSC-TSC differentiation to STB and induces potent innate immune response

We next investigated global gene transcriptional changes in infected eSTBs (Figure 4A). Pearson correlation and cluster analysis revealed the SARS-CoV-2 infected cells (24 hours post infection - hpi, or STB-D3; 48hpi or STB-D4) were clustered together and separated from the mock infection controls (Figure S4A). Comparative transcriptomic analysis identified 654 upregulated differentially expressed genes (DEGs) and 736 downregulated DEGs in SARS-CoV-2 infected cells (Figures S4B).

**Figure 4.**
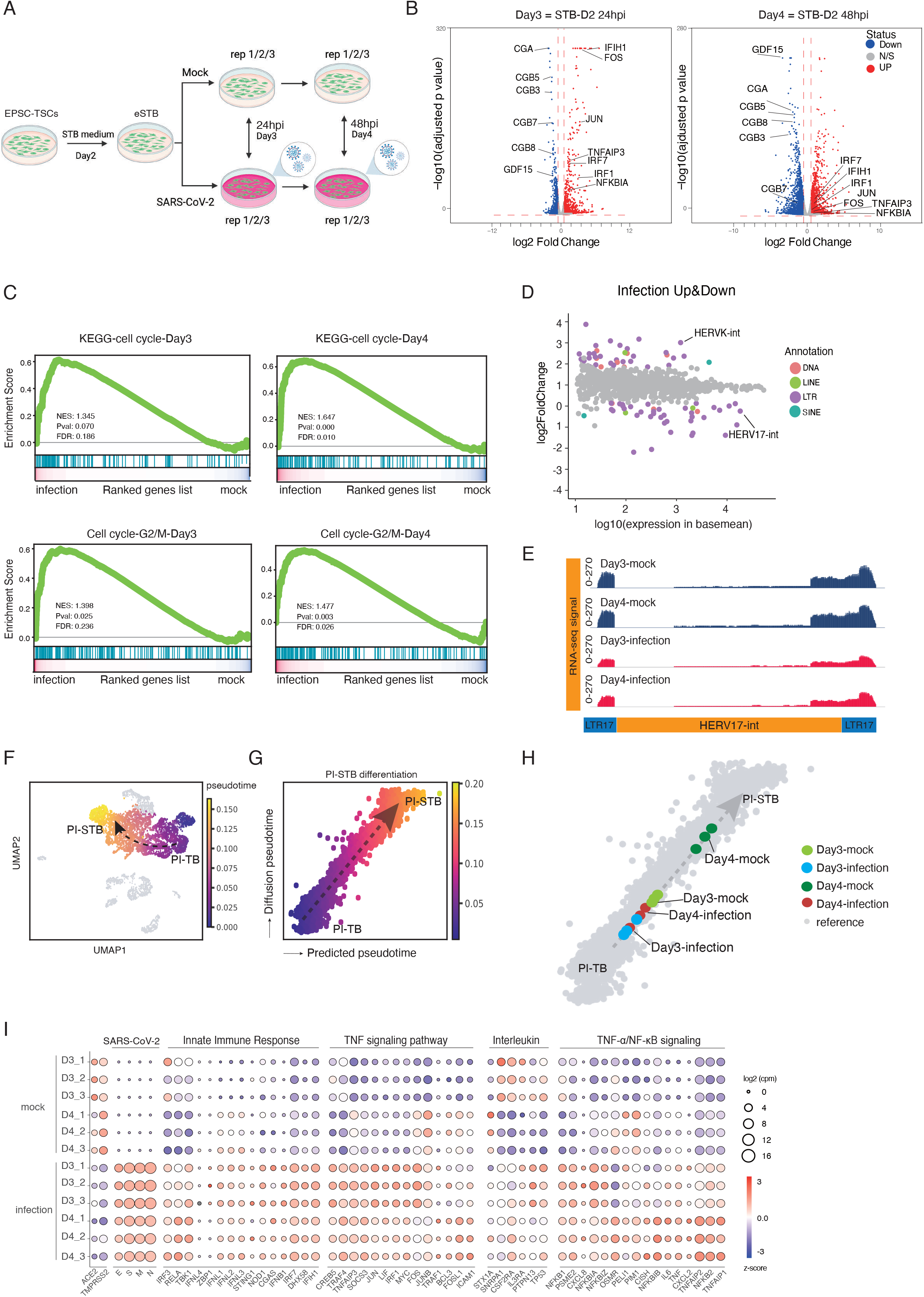
SARS-CoV-2 infection impairs differentiation toward STBs. (A) Experimental flow of eSTB generation for SARS-CoV-2 infection and sample collection for RNAseq. eSTBs used here are STB-D2. Samples were collected at 24 hpi (STB-D3 infection) and 48 hpi (STB-D4 infection) with three replicates. (B) Volcano plots of gene expression in infected STB-D3 and STB-D4. Significantly upregulated genes are labelled in red and significantly downregulated genes are labelled in blue. Horizontal red dash line marks adjusted P-value (Wald test) 0.05 and vertical lines marks expression fold change of 1.5. (C) Gene set enrichment analysis (GSEA) for cell cycle related genes including those of G2-M transition in virus infected STB-D3 and STB-D4 cells. Red lines are upregulated genes, blue lines are downregulated genes. (D) A scatter diagram of transcriptomic analysis of transposable elements (TEs) after virus infection. Differentially expressed TEs (P < 0.05, Wald test) are labelled with colours while none-differentially expressed TEs are labelled with grey colour. Expression levels of TEs used for x-axis are from DESeq2 results. Data of STB-D3 and STB-D4 are combined in this analysis. (E) RNAseq signal of HERV-W in infected STB-3 and STB-4 and the mock infection control cells. Library size was used to normalize the read counts. (F) Pseudotime analysis depicting PI-TB to PI-STB development trajectory. Dark purple indicates earlier pseudotime, light yellow indicates STB development. PI-TB to PI-STB pseudotime highlighted in the merged *in vivo* scRNA datasets as shown in Fig. 2B. Black dashed arrow indicates the imputed direction of differentiation. (G) PI-TB to PI-STB subpopulations colored by machine learning predicted pseudotime. X-axis is the predicted pseudotime, and y-axis is diffusion pseudotime computed using SCANPY. Grey dashed arrow describes the linear regression relationship between the predicted pseudotime and the diffusion pseudotime. (H) Infected vs. normal (mock) STB-D3 and STB-D4 mapped against the PI-TB to PI-STB pseudotime trajectory. STB cells are colored by differentiation time and treatment: light green for STB-D3 normal cells, blue for infected STB-D3, dark green for normal STB-D4, and red for infected STB-D4. The *in vivo* trophoblast reference is in grey. X-axis is the predicted pseudotime. For *in vivo* trophoblast, y-axis is diffusion pseudotime computed using SCANPY, whereas for the *in vitro* cells y-axis is the imputed observed pseudotime values imputed by *in vivo* linear regression equation (See Methods). Predicted pseudotime increases from bottom left to top right, as shown in Fig. 4G. (I) Bubble plot for ACE2, TMPRSS2, genes of SARS-CoV-2, innate immune response, TNF signaling pathway, interleukin signaling pathway and TNF-α/NF-κB signaling. Label color is according to z-score and bubble sizes are proportional to expression levels.

Among the genes that were significantly decreased in the SARS-CoV-2 infected cells were CGA, CGBs (CGB3, 5, 7, 8), GCM1, SDC1 and ENDOU and others that are highly expressed in mature and multinucleated STBs (Figure 4B and Figure S4C), indicating that the infected caused a possible developmental block. GSEA indicated a significant enrichment of cell cycle in particular G2-M genes in the infected cells compared to the mock control (higher cell cycle gene levels in the infected cells) (Figure 4C). The lack of mature STB signature and higher cell cycle genes in infected cells are consistent since mature and multinucleated STBs are known to have little cell cycle activities. On the other hand, SARS-CoV-2 infection did not substantially affect the infected cell viability as apoptosis-related genes were not significantly changed in the infected cells (Figure S4D).

eSTBs fuse and form multinucleate STBs, which depends on endogenous retrovirus (HERV) proteins Syncytin-1, an envelope gene of HERV-W, and Syncytin-2 produced from HERV-FRD. SARS-CoV-2 infection of eSTBs resulted in lower levels of both HERV-W (HERV17-int, Syncytin-1) and HERV-FRD (Syncytin-2) but substantially increased HERV-K expression (Figure 4D-4E, and Figures S4E). HERV-K is known to be exclusively expressed in cytotrophoblast cells (progenitors) in the human placenta (Kämmerer et al., 2011). These data indicate that SARS-CoV-2 infection impaired early STB differentiation and maturation.

To further investigate STB development, we next compared the transcriptomic pseudotime of the mock and infected STBs. EPSC-TSCs transcriptomically resembled PI-TB, and TSC-derived STB to PI-STB as described above in Figure 2C. We subtracted the PI-TB and PI-STB differentiation process from the *in vivo* trophoblast scRNAseq data (Figure 4F). The computed pseudotime increased from the immature PI-TB to the relatively mature PI-STB (Figures 4G). Using machine learning methods (validated as in Figure S4F), we mapped the mock and virus-infected STB cells (STB-D3) and STB-D4) to the PI-TB to PI-STB pseudotime trajectory, and discovered that the infected cells were closer to PI-TB than to the mock control cells along the pseudotime trajectory (Figure 4H). This result again demonstrates that SARS-CoV-2 infection of eSTBs adversely affects normal trophoblast development and function, which could be relevant to the reported early pregnancy miscarriages in some COVID-19 patients.

SARS-CoV-2 infection caused a strong innate response. Genes encoding interferon signaling components (IFNL1, IFIH1) and genes associated with TNFα signaling via NFKB such as TNFAIP3 and NFKBIA were all up regulated in the infected cells (Figures 4B and S4C). Gene Ontology (GO) term analysis found enriched GO terms related to virus cellular response and immune response pathways in SARS-CoV-2 infected STBs (Figure S4G). As expected, KEGG pathway analysis revealed that the coronavirus disease-COVID19 pathway was among the over-represented ones (Figure S4G). Bubble plot analysis further demonstrated in the upregulated genes an enrichment of those associated with innate immune response, interleukin and TNF signaling pathways (Figure 4I).

The double-stranded RNA (dsRNA), which is generated during coronavirus genome replication and transcription, could be recognized by melanoma differentiation gene 5 (MDA5/ IFIH1) in the cytoplasm to trigger innate immune activation upon coronavirus infection (Kindler et al., 2016; Li et al., 2010; Yin et al., 2021). IFIH1 (MDA5) was highly up regulated in the infected cells (Figure 4I). DNA is not known to be involved in the SARS-CoV-2 life cycle. Genes encoding cGAS and STING1, both being components of the cGAS-STING pathway of the innate immune system detecting cytosolic DNA, were not substantially altered (Figure 4I).

Type III IFNs have important antiviral functions (Ye et al., 2019). Particularly, IFN-λ1 is known to be constitutively released from human placental trophoblasts to protect the foetus from viral infection (Bayer et al., 2016; Chen et al., 2017). Type III IFNs have also been shown to restrict SARS-CoV-2 infection in airway and intestinal epithelia (Stanifer et al., 2020; Vanderheiden et al., 2020). Infected cells expressed high levels of Type III IFNs including IFN-λ1 (IL-29 or IFNL1), IFN-λ2 (IL-28A or IFNL2), IFN-λ3 (IL-28B or IFNL3) and IFN-λ4 (IFNL4) (Figure 4I).

### Remdesivir and GC376 effectively eliminate eSTB SARS-CoV-2 infection

eSTBs permitted robust SARS-CoV-2 infection and thus provide normal and physiologically relevant cells for evaluating anti-SARS-CoV-2 drugs. To this end, we tested the FDA-approved antiviral remdesivir (Rubin et al., 2020) and a veterinary drug GC376 (Ma et al., 2020) (Figure 5A). Remdesivir effectively eliminated SARS-CoV-2 infection in Vero E6 cells at around 5μM (Wang et al., 2020). Remarkably, in eSTBs, as low as 0.08μM of remdesivir decreased SARS-CoV-2 viral load by three orders of magnitudes (Figure 5B). Similar antiviral potency of remdesivir treatment was observed in eSTBs infected by MERS-CoV, another highly pathogenic human coronavirus (Figure 5C). GC376 is a repurposed SARS-CoV-2 main protease inhibitor that can increase survival of mice with a fatal SARS-CoV-2 infection (Dampalla et al., 2021). GC376 suppressed both SARS-CoV-2 and MERS-CoV in eSTBs (Figures 5D and 5E). The effectiveness of these two drugs against SARS-CoV-2 was further confirmed by the remarkably reduced viral N antigen expression in immunofluorescence staining (Figure S5A), and by the substantially reduced SARS-CoV-2- and MERS-CoV-induced cytopathogenic effects (CPE) (Figures S5B-S5E).

**Figure 5.**
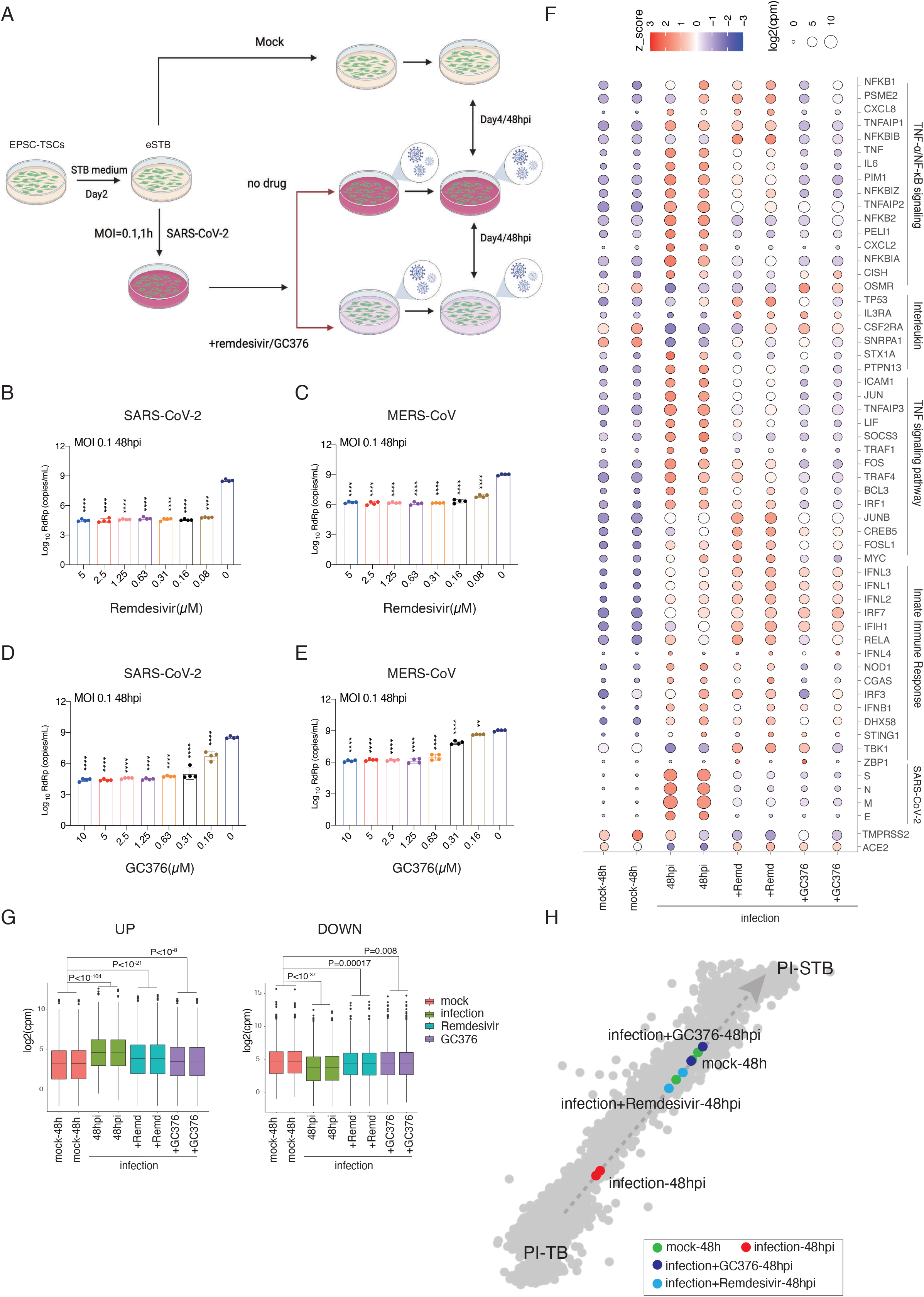
Low dosages of antiviral drugs effectively eliminated SARS-CoV-2 and MERS-CoV infection of eSTBs. (A) Experimental design for drug treatment. eSTBs (STB-D2) were submitted for SARS-CoV-2 or MERS-CoV infection at 0.1 MOI for one hour followed by incubation with or without remdesivir and GC376. Cells were collected at 48 hpi for analysis. (B-C) RT-qPCR analysis of virus genome copy in the cell culture supernatant of 48 hpi eSTBs treated with remdesivir. (B) (SARS-CoV-2); (C) MERS-CoV. Values are mean ± SEM from three independent experiments. *** p < 0.001 (Ordinary one-way ANOVA). (D-E) RT-qPCR analysis of virus genome copy in the cell culture supernatant of 48hpi eSTBs treated with GC376. (D) (SARS-CoV-2); (E) MERS-CoV. Values are mean ± SEM from 3 separate experiments. *** p < 0.01, *** p < 0.001 (Ordinary one-way ANOVA). (F) Bubble plot for genes of TNF-α/NF-κB signaling, interleukin signaling pathway, TNF signaling pathway, innate immune response and SARS-CoV-2. Color is according to expression z-score and bubble size is expression level. (G) Box plots for up-regulated and down-regulated genes in 48 hpi eSTBs, with or without SARS-CoV-2 infection, and in the presence or absence of remdesivir or GC376. Statistical significance was calculated by Wilcoxon test using mean of expression value. (H) Mapping of 48 hpi eSTBs (in the presence or absence of remdesivir or GC376) against in-vivo pseudotime trajectory. Pseudotime increases from bottom left to top right, as shown in Fig 4G. The cells with different treatment were colored: red for infected cells without any treatment, green for mock (non-infected cells, no drug treatment), light blue for infected cells treated by remdesivir, and dark blue for infected cells treated by GC376.

We next performed global gene expression analysis of the infected and drug-treated cells. Both drug treatments drastically reduced expression of SARS-CoV-2 genes (E, M, N, and S) (Figure 5F). Neither one appeared to cause substantial changes of host innate immune response genes such as MDA5, IFN-λs and IFNb. On the other hand, proinflammatory cytokine genes such as TNF, IL-6 and IL-8 were all down-regulated in the drug treatment cells (Figures 5F and S5F).

SARS-CoV-2 infection of eSTBs impaired STB differentiation as presented in Figure 4. We assessed whether remdesivir and GC376 treatment and mitigation of infection could rescue the development. Hierarchical clustering analysis confirmed a shift towards the mock infection control STBs (Figure S5G). The global up-regulated and down-regulated genes in the infected-drug-treated cells were comparable to that in the mock infected cells (Figure 5G). Furthermore, global expression analysis revealed that along the PI-TB to PI-STB pseudotime trajectory, remdesivir and GC376 treatment of the infected STB cells caused a shift away from PI-TB and to PI-STB, indicating that antiviral treatment drugs partially rescued STB development and differentiation (Figure 5H).

### Direct derivation of 3D trophoblast organoids from EPSC-TSCs for SARS-CoV-2 infection

Long term and genetically stable trophoblast organoids could be derived from first trimester placenta tissues or the blastocyst, which grow as complex structures that closely recapitulate the organization of *in vivo* placental villi (Haider et al., 2018; Turco et al., 2018). It was recently reported that 3D or organoid culture of hTSCs more closely resembled the *in vivo* counterparts than 2D hTSCs (Sheridan et al., 2021). The observation that eSTBs were highly susceptible to SARS-CoV-2 prompted us to investigate whether the 3D trophoblast organoids could be adopted for studying SARS-CoV-2 infection. From EPSC-TSCs, we directly established trophoblast organoids under a published culture condition (TOM) (Turco et al., 2018), where individual hTSCs self-aggregated into small clumps in a few days and eventually matured into 3D organoids in about 20 days (Figure 6A). The organoids were mechanically dissociated, passaged, and maintained for at least 10 passages. hTSC-derived organoids organized into villous-like structures which were reminiscent of those derived from the placenta (Haider et al., 2018; Turco et al., 2018), where the basement membrane was on the outside in contact with the Matrigel substratum, whereas syncytial masses lined the central cavity (Figure 6B). Multinucleated mature STBs expressing CGB and ENDON were found inside the organoids and did not express the stemness transcription factors GATA3, TFAP2A and TFAP2C or eSTB marker CD46 (Figures 6C, and Figures S6A and S6B). Trophoblast organoids harbored both stem cells and STBs (Haider et al., 2018; Turco et al., 2018). They expressed genes of both hTSCs and STBs including GATA3, ITGA6 and TEAD4, and ERVW-1 and CGB, although the levels were lower compared to those in hTSCs or pure STB cultures, respectively (Figure 6D, and Figure S6C). Nevertheless, ELISA detected full-length and properly folded β-hCG hormone secreted by trophoblast organoids (Figure S6D). Consistent with a recent study (Sheridan et al., 2021), trophoblast organoids expressed higher levels of some trophoblast-specific miRNAs (has-miR-517a and 525-3p) compared to the 2D EPSC-TSCs (Figure S6E). Although trophoblast organoids did not express appreciable levels of EVT genes, they robustly generated migrating HLA-G^+^ and ITGA5^+^ EVT cells (Figure S6F-S6G), confirming the presence of bi-potential stem cells.

**Figure 6.**
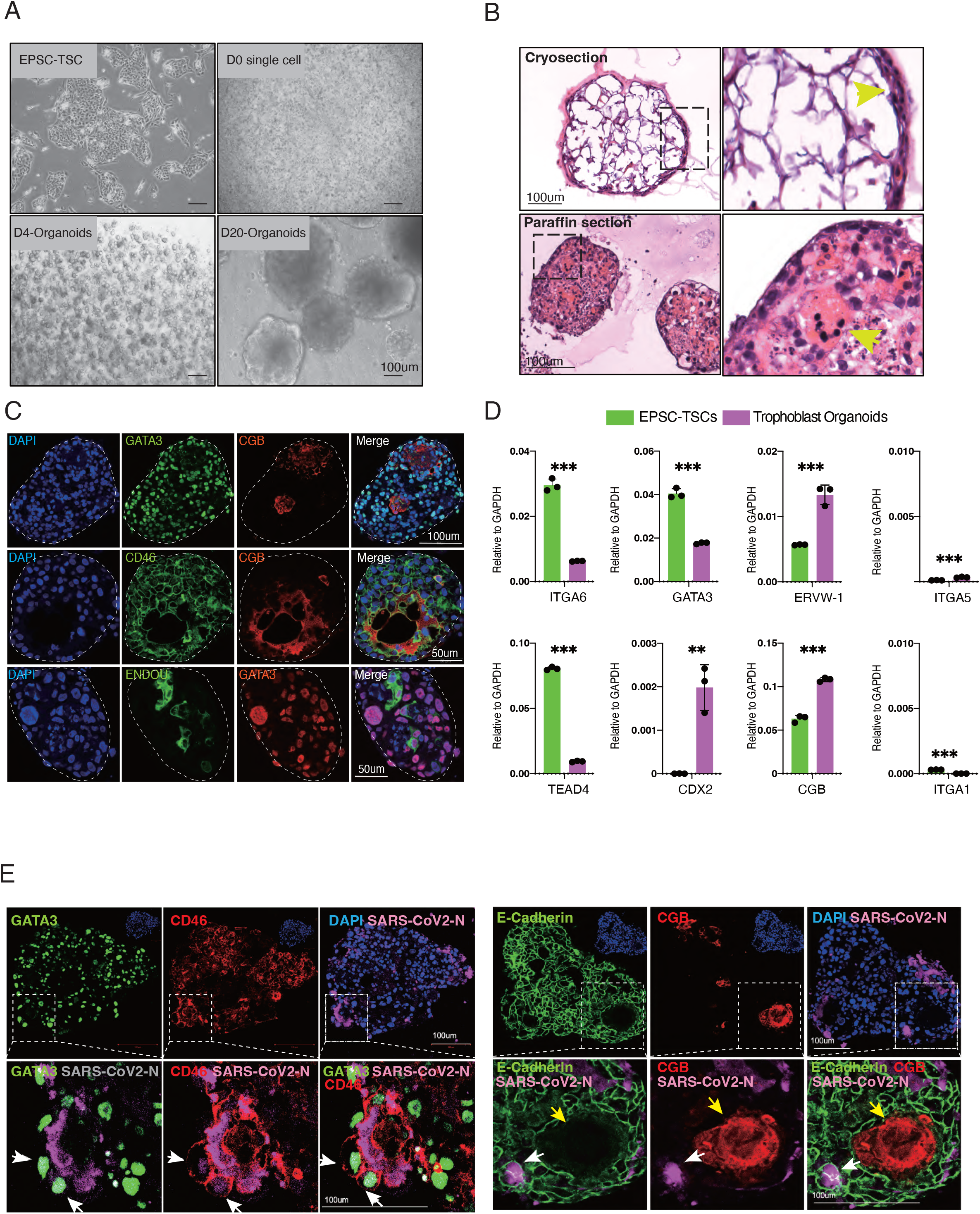
Infection of EPSC-TSC trophoblast organoids by SARS-CoV-2. (A) Generation of trophoblast organoids from EPSC-TSCs. Morphology of single cell dissociated EPSC-TSCs (day 0), and cell clumps on day 4 and trophoblast organoids on day 20. Scale bar: 100μm (B) H&E staining of cryosections (upper panels) and paraffin sections (lower panels) of trophoblast organoids. Cells at the organoid peripheral are progenitors (Haider et al., 2018; Turco et al., 2018). Arrowhead in the cryosection (upper panel) indicates densely packed cell clusters in the outer layer of trophoblast organoids. Arrowhead in the paraffin section (lower panel) indicates the multinucleated STBs inside the trophoblast organoid. Scale bar: 100μm (C) Immunofluorescence staining of trophoblast organoid cryosections for GATA3 and for STB markers CD46, CGB and ENDOU. DAPI stains the nucleus. Scale bar: top panel 100μm; mid and lower panels 50μm (D) RT-qPCR analysis of trophoblast transcription factor genes (TEAD4, CDX2, ITGA6 and GATA3), STB genes (ERVW-1 and CGB) and EVT genes (ITGA1 and ITGA5) in EPSC-TSCs and trophoblast organoids (Data are mean ± SD, n=3 independent replicates from 3 separate sample extracts. ** p < 0.01; *** p < 0.001 (two-tailed unpaired Student’s t-test). Gene expression levels were normalized to GAPDH using the ΔCt method. (**E**) Immunofluorescence staining paraffin sections of SARS-CoV-2 infected trophoblast organoids for GATA3, CD46, E-Cadherin and CGB. Most infected cells were stained positive for CD46, but were low or negative for CGB. White arrows indicate the mononucleated infected cells, whereas yellow arrows point to a large multinucleated STB which expresses high CGB. The dash line box areas are shown in higher resolution. Nuclei were counterstained by DAPI. Scale bars: 100μm

We next infected trophoblast organoids by co-culturing with SARS-CoV-2 at an MOI of 10 in TOM and harvested them after 48 hours infection for analysis. SARS-CoV-2 N protein was found in virus co-cultured organoid cells that expressed CD46, but not in those multinucleated cells with high CGB expression (Figures 6E and S6H), in line with the 2D culture results that eSTBs or more immature STBs were the ones infected. Trophoblast organoids could therefore provide a novel 3D tissue architecture for studying SARS-CoV-2 infection, which could be further explored by including immune cells in the organoids.

### ACE2 is essential for SARS-CoV-2 infection in trophoblasts

In the peri-implantation embryos, PI-STBs were the only trophoblasts that co-expressed ACE2 and TMPRSS2 (Figure 2E). High levels of ACE2 protein were detected in eSTBs (Figure 3F), and all the SARS-CoV-2 infected trophoblasts expressed ACE2 (Figure 3C, 3K and Figure S3G). To genetically examine the role of ACE2 in SARS-CoV-2 infection in human trophoblasts, we knocked out ACE2 in hEPSCs using the CRISPR/Cas9 and made homozygous deletions in coding exon 2 (Figures S7A-S7D). The ACE2 knockout hEPSCs (ACE2-KO) had normal morphology and expressed high levels of key pluripotency factors (OCT4 and NANOG) and markers (SSEA4 and TRA-1-60) but low levels of lineage genes (CDX2, ELF5, SOX17, GATA6, SOX1) (Figures 7A and S7E-7F), comparable to the normal parental hEPSCs.

**Figure 7.**
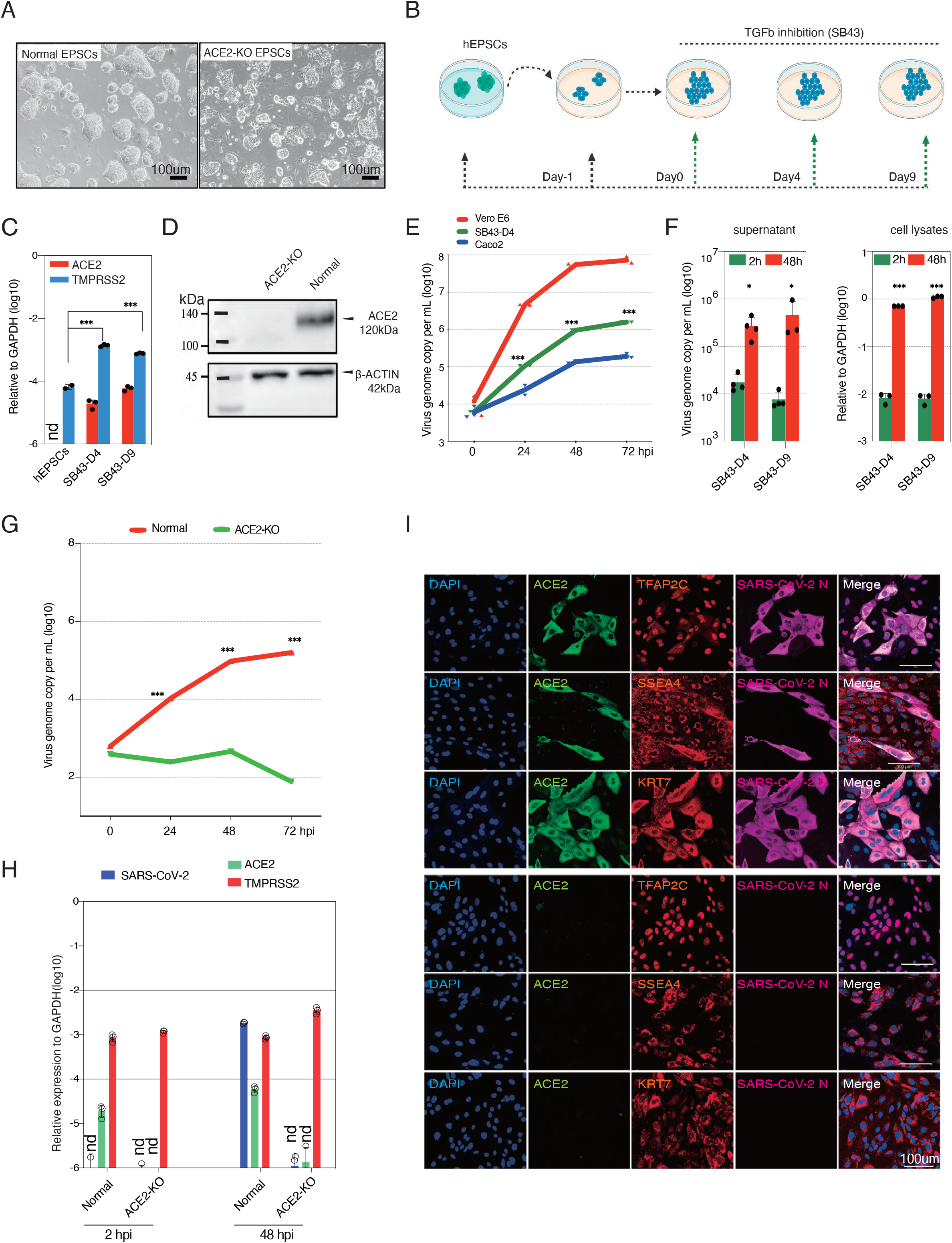
ACE2 is required for SARS-CoV-2 infection of trophoblasts differentiated from hEPSCs. (A) Normal and ACE2-KO hEPSCs. Scale bar: 100μm (B) Experimental flow of hEPSCs differentiation towards trophoblasts by the TGF-β inhibitor SB-431542 (SB43) treatment. This method directly produces trophoblasts, primarily STBs, from hEPSCs. Experiment details are in Methods. (C) RT-qPCR analysis of genes encoding ACE2 and TMPRSS2 in hEPSCs and the differentiated cells on day 4 and day 9. hEPSCs have no ACE2 expression. nd: undetectable. (Data are mean ± SD, n=3 independent replicates from 3 separate sample extracts. ** p < 0.01; *** p < 0.001 (two-tailed unpaired Student’s t-test). Gene expression was normalized to GAPDH using the ΔCt method. (D) Western blotting confirms loss of ACE2 protein in the SB43-treated hEPSCs on day 9 (SB43-D9) of differentiation. (E) SARS-CoV-2 replication kinetics in cells differentiated from hEPSCs compared to Vero E6 and Caco2 cells. Caco2 cell, an immortalized cell line derived from human colorectal adenocarcinoma and is often used in infection studies of SARS-CoV-2. All cells were infected with SARS-CoV-2 at 0.1MOI. Viral supernatant samples were harvested at 2, 24, 48 and 72 hpi for RT-qPCR of viral RNA load. (Data are mean ± SD, n=3 independent replicates from 3 separate experiments. *** p < 0.001 (two-tailed unpaired Student’s t-test). (F) Left panel: RT-qPCR of supernatant viral load in SB43-treated hEPSCs at 2 and 48 hpi. Cells were challenged with SARS-CoV-2 at 0.1 MOI. (Data are mean ± SD, n=3 independent replicates from 3 separate experiments. * p < 0.05 (two-tailed unpaired Student’s t-test). Right panel: RT-qPCR quantification of SARS-CoV-2 genome in cell lysates of SB43-treated hEPSCs on days 4 and day 9. (Data are mean ± SD, n=3 independent replicates from 3 separate sample extracts. *** p < 0.001 (two-tailed unpaired Student’s t-test). Gene expression was normalized to GAPDH using the ΔCt method. (G) Quantification of supernatant viral RNA loads of SB43-treated (day 4) normal and ACE2-KO EPSCs. Cells were infected with SARS-CoV-2 at 0.1MOI and supernatant samples were harvested at 2, 24, 48 and 72 hpi. (Data are mean ± SD, n=3 independent replicates from 3 separate experiments. *** p < 0.001 (two-tailed unpaired Student’s t-test). (H) SARS-CoV-2 viral genome quantitation and expression of ACE2 and TMPRSS2 in normal versus SB43-treated ACE2-KO EPSCs at day 4. (Data are mean ± SD, n=3 independent replicates from 3 separate sample extracts. nd, not detectable. Gene expression was normalized to GAPDH using the ΔCt method. (I) Representative immunofluorescence staining images for ACE2, trophoblast factors and markers TFAP2C, SSEA4, and KRT7, and SARS-CoV-2 N protein in normal (upper panels) and ACE2-KO cells (lower panels) (day4 of SB43-treatment) at 48 hpi. No SARS-CoV-2 N protein was detected in the ACE2-KO cells. Scale bars: 100μm

**Figure 8.**
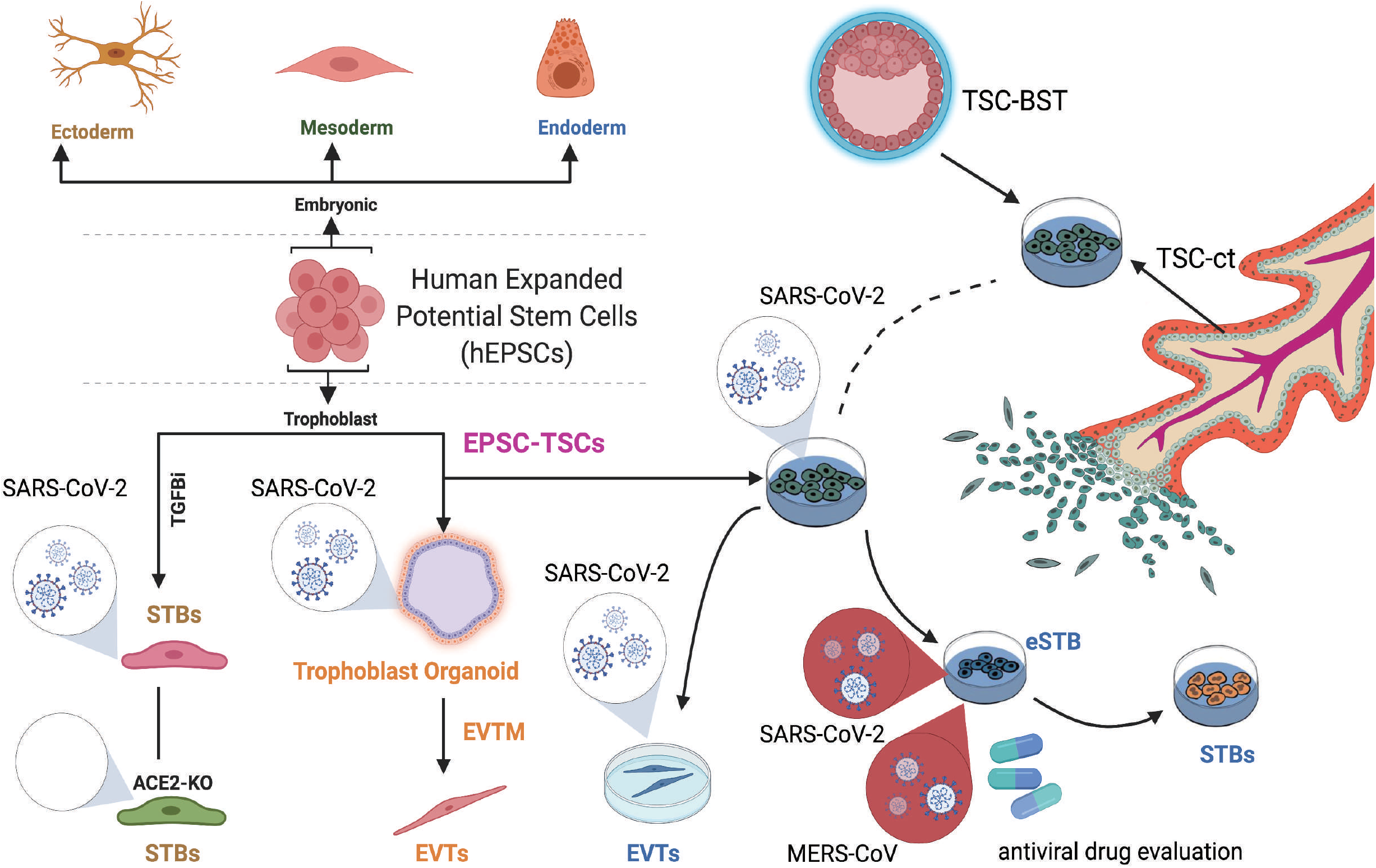
Schematic of studying human EPSC-derived or placenta/blastocyst-derived trophoblast susceptibility to SARS-CoV-2 and MERS-CoV. Human EPSCs retain developmental potential to all cell types of both embryonic and extra-embryonic cell lineages including trophoblasts. With TGF-β inhibition, human EPSCs can directly differentiate to trophoblasts, which can be infected with SARS-CoV-2. Knocking out ACE2, which mediates SARS-CoV-2 recognition and membrane fusion, makes SARS-CoV-2 unable to infect trophoblasts. TSCs can be established from hEPSCs, or from human blastocysts (TSC-BST) or placenta cytotrophoblast (TSC-ct). EPSC-TSCs can differentiate to trophoblast subtypes STBs and EVTs. EPSC-TSCs can also generate 3D trophoblast organoids, which recapitulate the physiological features of trophoblast organoids directly derived from the human placenta. eSTBs (early STBs) have high ACE2 expression and are highly susceptible to SARS-CoV-2 infection. eSTBs can serve as a new cell model for coronavirus research and for antiviral drug evaluation.

We next induced normal and ACE2-KO hEPSCs to trophoblasts using a simple and efficient protocol by the TGF-β inhibitor SB431542 (Gao et al., 2019) (Figures 7B and S7G). Normal hEPSCs did not express ACE2 (Figures 7C) and were not infected by SARS-CoV-2 as described above (S3A and S3B). The differentiated cells expressed ACE2, TMPRSS2, and typical STB markers such as SDC1, ERVW-1 (Syncytin-1), GATA3, KRT7 and CGB (Figure 7C, and Figures S7H and S7I). In contrast, no ACE2 protein was detected in cells differentiated from the ACE2-KO hEPSCs (Figure 7D).

The cells differentiated from normal and ACE2-KO hEPSCs were subsequently subjected to SARS-CoV-2 infection. We selected cells of day-4 and day-9 differentiation for infection as they expressed ACE2 and TMPRSS2 (Figure 7C). Infection of these cells produced substantial amounts of viral genome in the supernatant and cell lysates (Figures 7E and 7F). The infected cells, detected as SARS-CoV-2 nucleocapsid (N) protein-positive, all expressed ACE2 and trophoblast factor and markers TFAP2C, SSEA4 and KRT7, and most were mononucleated (Figure 7I and S7J). Loss of ACE2 abolished the infection as indicated by the drastic decrease of viral genome copy numbers in the supernatants (Figure 7G) and in cell lysates (Figure 7H). In immunofluorescence staining, no cells were stained positive for either SARS-CoV-2 N protein or ACE2 in the cells differentiated from ACE2-KO EPSCs (Figure 7I). Therefore, in human trophoblasts, ACE2 is essential for SARS-CoV-2 infection.

## Discussion

Our work presented here provides proof-of-concept results that human EPSCs can provide a cellular system for assessing susceptibility of trophoblasts to SARS-CoV-2 and other coronavirus infection at high resolution. Among the cell types examined including hEPSCs, hTSCs, STBs and EVTs, eSTBs are highly susceptible to SARS-CoV-2 infection in both 2D and 3D models. This discovery was made possible by the hEPSC-based *in vitro* system since eSTBs are only transiently present in the placenta trophoblast development and the non-proliferative multinucleated STBs quickly form the layer covering placenta villi. Neither trophoblastic cell lines nor primary trophoblasts might be representative of these eSTBs. Importantly, our findings are in line with the clinical reports that opportunistic SARS-CoV-2 infection during pregnancy can occur but is an infrequent event. The eSTB susceptibility to SARS-CoV-2 may implicate a higher risk in early pregnancy. hEPSCs have the developmental potential to generate all embryonic and extraembryonic cell lineages. Besides trophoblasts, hEPSCs can be explored to derive diverse cell types and to functionally assess their susceptibility to SARS-CoV-2, including airway epithelial, gastrointestinal epithelial and neuronal cells. hEPSCs are robust in culture and permit efficient precision genome editing. These qualities grant them potentials in overcoming some of the long-standing technical challenges such as cell transfection and genetic manipulation of various target cells. VeroE6 cells originated from frican green monkey are commonly used for isolation and propagation of SARS-CoV-2 (Zhou et al., 2020) and have been approved for use in vaccine manufacturing. VeroE6 have genetic defects including large genomic deletions encompassing the interferon gene clusters (IFN-a and -b) and the CDKN2A/B loci and are thus deficiencies in IFN response (Osada et al., 2014), which have prevented its application in the study of antiviral response. Furthermore, the lack of TMPRSS2 in VeroE6 cells results in the selection of SARS-CoV-2 variants that have lost the polybasic furin cleavage site at the S1-S2 junction (Sasaki et al., 2021). It is thought that SARS-CoV-2 enters VeroE6 cells mainly through the less specific clathrin-mediated endocytic pathway but cannot use the TMPRSS2-facilitated pathway based on the binding of S with ACE2. In contrast, hESPCs are normal human cells with an intact innate immune system but without major known genetic or epigenetic defects.

EPSC-derived eSTBs are capable of using ACE2 and TMPRSS2 for SARS-CoV-2 entry into cells and are highly potent to support SARS-CoV-2 propagation. Hence, hEPSC-derived cells have the potential to be developed as new virus producer cells and inspire novel cell-based therapeutics for various infections. Finally, hEPSC-derived organoids, particularly those harboring immune cells, can potentially provide a more physiologically relevant system to study pathogenesis and immunopathogenesis of SARS-CoV-2 and other pathogens.

## STAR*METHODS

Detailed methods include the following:

- **KEY RESOURCES TABLE**
- **RESOURCE AVAILABILITY**
  - Lead contact
  - Materials availability
  - Data and code availability
- **EXPERIMENTAL MODEL AND SUBJECT DETAILS**
- **METHODS DETAILS**
  - Culture of hEPSCs
  - Differentiation of hEPSCs to trophoblast lineages by the TGF-β inhibitor SB431542
  - Derivation of human Trophoblast Stem Cell (hTSCs) from hEPSCs.
  - Differentiation of EPSC-TSCs to STBs
  - Differentiation of EPSC-TSCs to EVTs
  - Generation of trophoblast organoids from EPSC-TSCs
  - Other human cell lines for SARS-CoV-2 infection
  - SARS-CoV-2 infection and detection
  - Detection of SARS-CoV-2 virus
  - Plaque assay
  - Guide RNA design, RNA synthesis and plasmid DNA preparation
  - Electroporation and selection
  - Genotyping and Sanger sequencing
  - Reverse Transcription-Quantitative Polymerase Chain Reaction (RT-qPCR)
  - Immunofluorescence staining
  - Western blotting
  - Flow cytometry
  - Transwell invasion assay
  - Statistical analysis and reproducibility
  - RNA-seq analysis
  - scRNA Preprocessing
  - Integration of in vitro and in vivo extra-embryonic datasets
  - Whole-transcriptome correlation analysis between in vitro cell line bulk RNA-seq and in vivo scRNA-seq
  - Integrative pseudotime analysis

- **QUANTIFICATION AND STATISTICAL ANALYSIS**

## SUPPLEMENTAL INFORMATION

SI Methods and Tables S1 to S3 include further details of the study materials and methods.

## ACKNOWLEDGEMENTS

This work was financially supported by National Key Research and Development Program of China (No. 2018YFA0902702), the National Natural Science Foundation of China (No. 81570202), Hong Kong Health and Medical Research Fund (No. COVID190114) and Hong Kong Research Grants Council (No. C7142-20GF), China Postdoctoral Science Foundation (2021M702280), High Level-Hospital Program, Health Commission of Guangdong Province, China (No. HKUSZH201902025), and the HKU internal funding (P. L.)

## AUTHOR CONTRIBUTIONS

Pentao Liu, Fang Liu, Degong Ruan and Dong-Yan Jin conceived the project and designed the experiments. Fang Liu performed the SB43 differentiation related experiments and ACE2 knockout to establish mutant hEPSCs; Zi-Wei Ye and Shuofeng Yuan performed all infection studies including viral load analysis; Degong Ruan performed the hTSC differentiation, trophoblast organoid generation and characterization, immunofluorescence staining and FACS assays; Ka-ki Tam performed RT-qPCR, analyzed the data and helped with figure revision. The three co-first authors, Pentao Liu and Dong-Yan Jin wrote the manuscript together. Jilong Guo, Yiyi Xuan, Zhuoxuan Li contributed to the CRISPR technology, Western blotting and re-analysis of RNA-Seq data, respectively. Chon-Phin Ong, Shuofeng Yuan and Kaiming Tang helped with viral infection and RT-qPCR analysis of viral RNA load. Pentao Liu and Dong-Yan Jin contributed to the critical revisions of the manuscript. All authors read and approved the final manuscript.

