## Supplemental Figures 1-7 for "Human Early Syncytiotrophoblasts Are Highly Susceptible to SARS-CoV-2 Infection"

Supplemental Text and Figures

### Figure S1

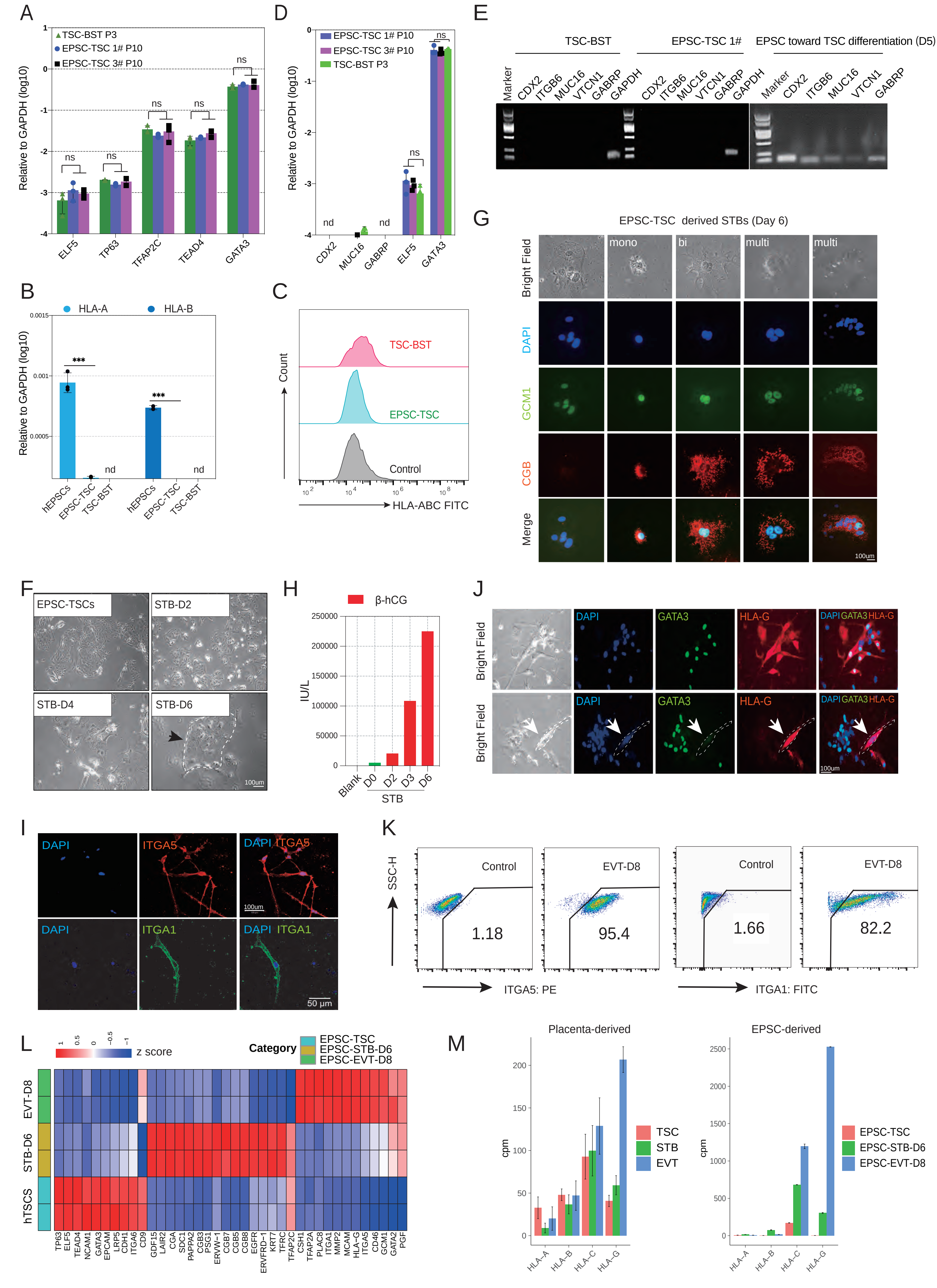

### Figure S2

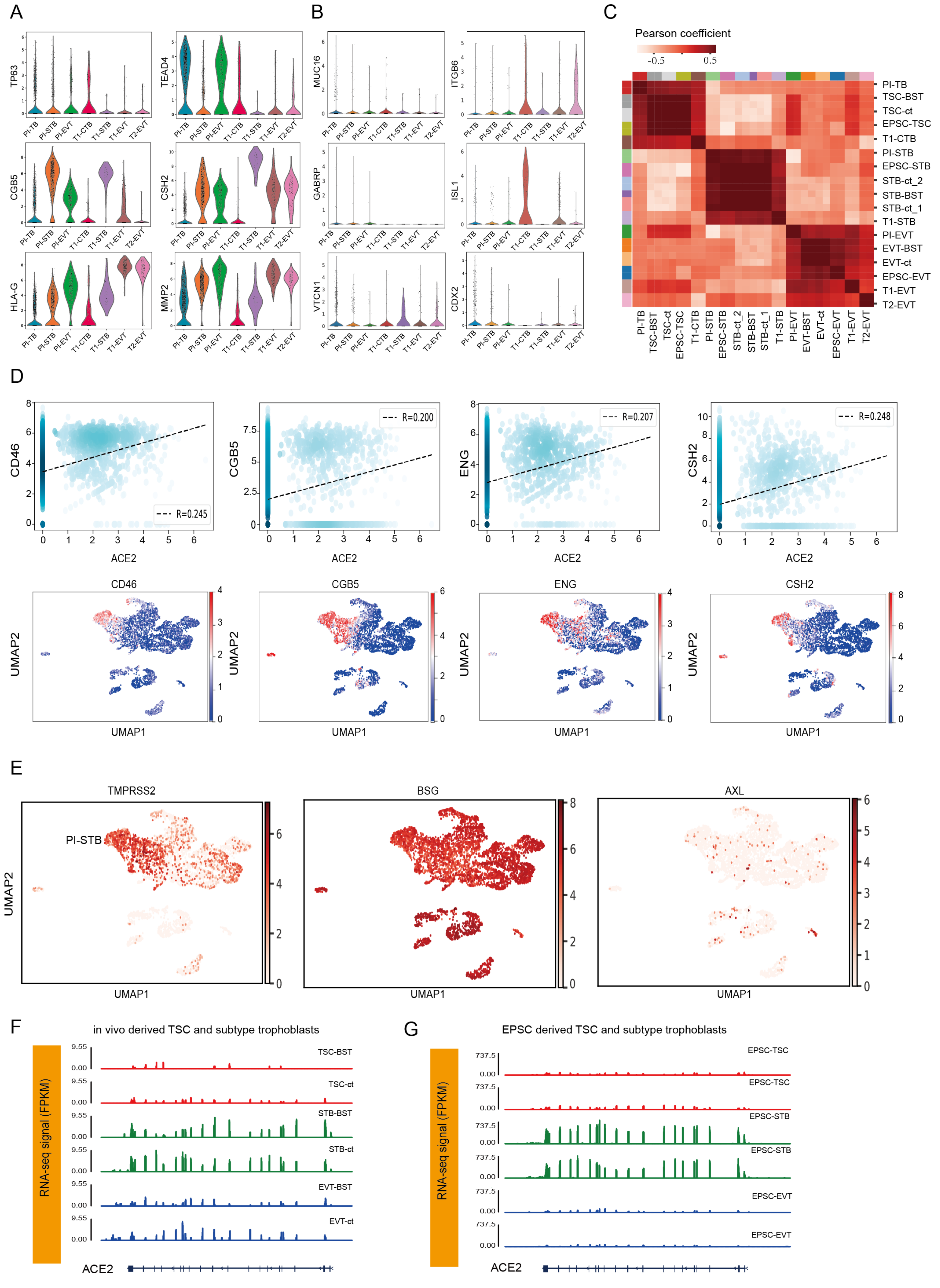
A
B
C
Pearson coefficient
-0.5
0.5
PI-TB
TSC-BST
TSC-ct
EPSC-TSC
T1-CTB
PI-STB
EPSC-STB
STB-ct_2
STB-BST
STB-ct_1
T1-STB
PI-EVT
EVT-BST
EVT-ct
EPSC-EVT
T1-EVT
T2-EVT
TP63
TEAD4
CGB5
CSH2
HLA-G
MMP2
MUC16
ITGB6
GABRP
ISL1
VTCN1
CDX2
D
CD46
CGB5
ENG
CSH2
ACE2
R=0.245
R=0.200
R=0.207
R=0.248
UMAP1
UMAP2
E
TMPRSS2
BSG
AXL
PI-STB
UMAP1
UMAP2
F
in vivo derived TSC and subtype trophoblasts
RNA-seq signal (FPKM)
TSC-BST
TSC-ct
STB-BST
STB-ct
EVT-BST
EVT-ct
9.55
0.00
ACE2
G
EPSC derived TSC and subtype trophoblasts
RNA-seq signal (FPKM)
EPSC-TSC
EPSC-STB
EPSC-EVT
737.5
0.00
ACE2

### Figure S3

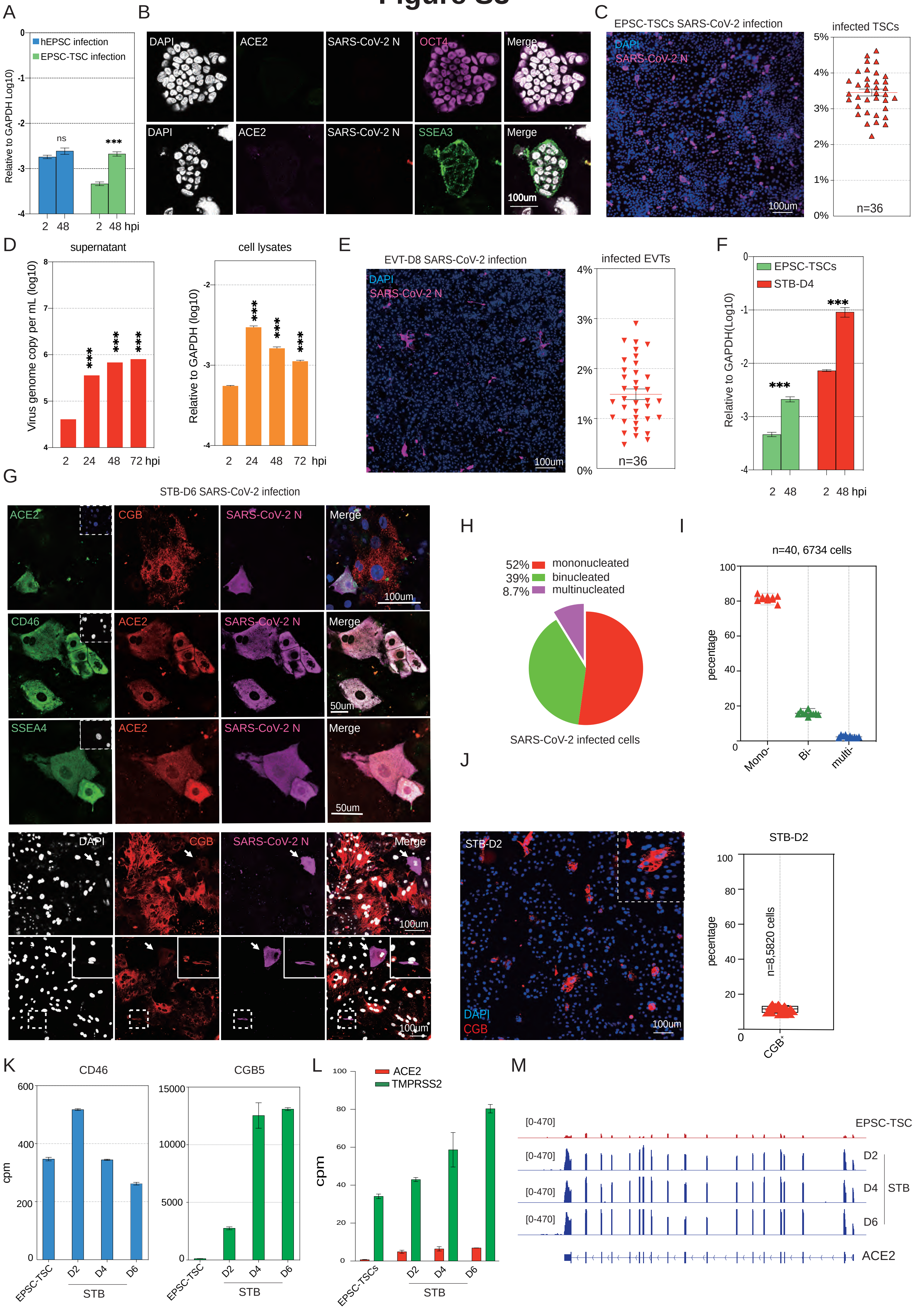
A
hEPSC infection
EPSC-TSC infection
Relative to GAPDH Log10)
ns
***
2 48 2 48 hpi
B
DAPI
ACE2
SARS-CoV-2 N
OCT4
Merge
SSEA3
100um
C
EPSC-TSCs SARS-CoV-2 infection
DAPI
SARS-CoV-2 N
100um
infected TSCs
n=36
D
supernatant
cell lysates
Virus genome copy per mL (log10)
Relative to GAPDH (log10)
2 24 48 72 hpi
E
EVT-D8 SARS-CoV-2 infection
infected EVTs
n=36
F
EPSC-TSCs
STB-D4
Relative to GAPDH(Log10)
G
STB-D6 SARS-CoV-2 infection
CGB
CD46
SSEA4
50um
H
52% mononucleated
39% binucleated
8.7% multinucleated
SARS-CoV-2 infected cells
I
n=40, 6734 cells
pecentage
Mono-
Bi-
multi-
J
STB-D2
CGB
n=8,5820 cells
CGB+
K
CD46
CGB5
cpm
EPSC-TSC
D2
D4
D6
STB
L
ACE2
TMPRSS2
EPSC-TSCs
M
[0-470]
EPSC-TSC
D2
D4
D6
STB
ACE2

### Figure S4

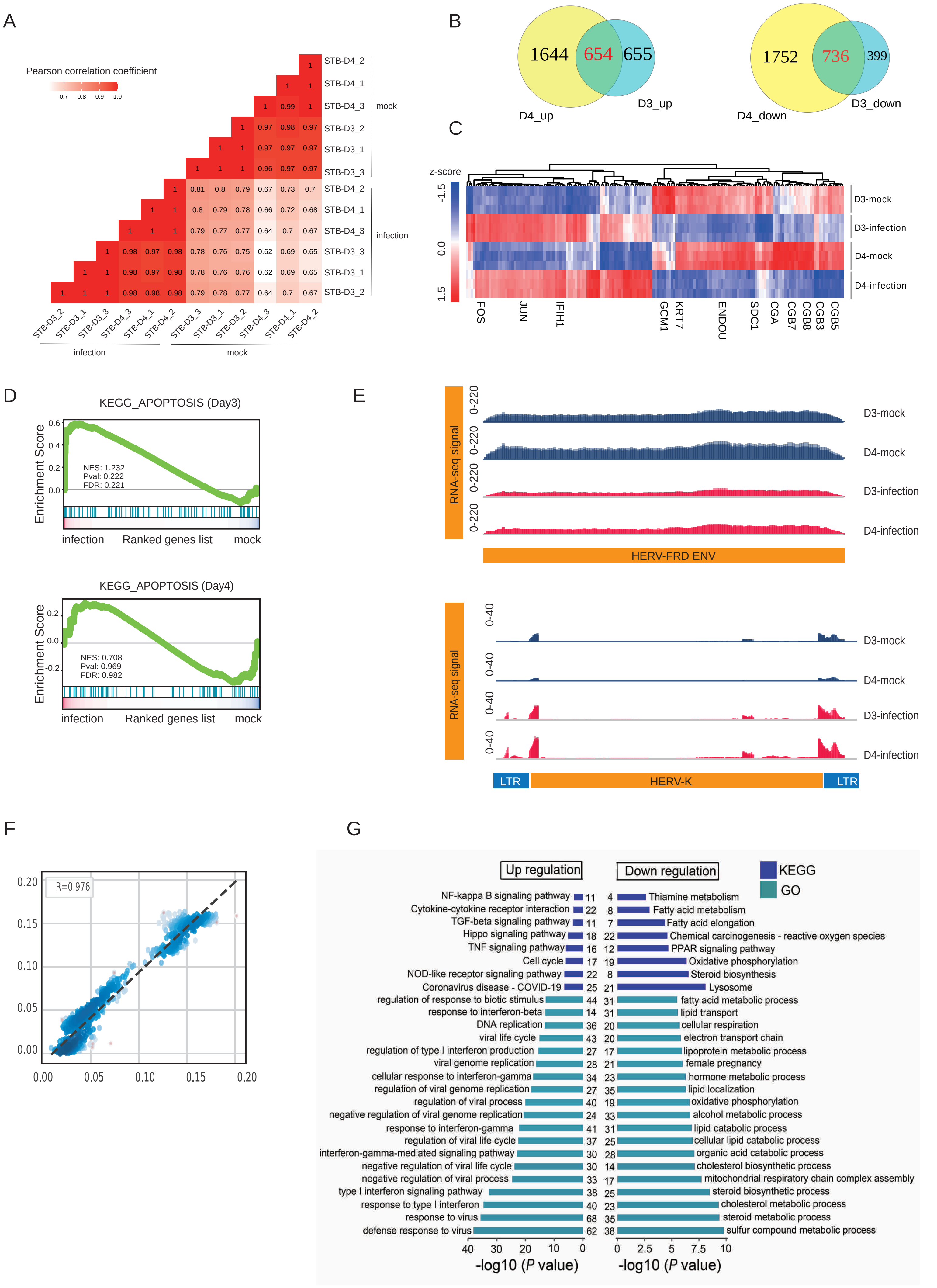

A
Pearson correlation coefficient
0.7 0.8 0.9 1.0
STB-D4_2
STB-D4_1
STB-D4_3
STB-D3_2
STB-D3_1
STB-D3_3
mock
infection
infection
mock
B
1644
654
655
D4_up
D3_up
1752
736
399
D4_down
D3_down
C
z-score
-1.5
0.0
1.5
D3-mock
D3-infection
D4-mock
D4-infection
FOS
JUN
IFIH1
GCM1
KRT7
ENDOU
SDC1
CGA
CGB7
CGB8
CGB3
CGB5
D
KEGG_APOPTOSIS (Day3)
Enrichment Score
NES: 1.232
Pval: 0.222
FDR: 0.221
infection
Ranked genes list
mock
KEGG_APOPTOSIS (Day4)
NES: 0.708
Pval: 0.969
FDR: 0.982
E
RNA-seq signal
0-220
D3-mock
D4-mock
D3-infection
D4-infection
HERV-FRD ENV
0-40
LTR
HERV-K
LTR
F
R=0.976
G
Up regulation
Down regulation
KEGG
GO
NF-kappa B signaling pathway 11
Cytokine-cytokine receptor interaction 22
TGF-beta signaling pathway 11
Hippo signaling pathway 18
TNF signaling pathway 16
Cell cycle 17
NOD-like receptor signaling pathway 22
Coronavirus disease - COVID-19 25
regulation of response to biotic stimulus 44
response to interferon-beta 14
DNA replication 36
viral life cycle 43
regulation of type I interferon production 27
viral genome replication 28
cellular response to interferon-gamma 34
regulation of viral genome replication 27
regulation of viral process 40
negative regulation of viral genome replication 24
response to interferon-gamma 41
regulation of viral life cycle 37
interferon-gamma-mediated signaling pathway 30
negative regulation of viral life cycle 30
negative regulation of viral process 33
type I interferon signaling pathway 38
response to type I interferon 40
response to virus 68
defense response to virus 62
4 Thiamine metabolism
8 Fatty acid metabolism
7 Fatty acid elongation
22 Chemical carcinogenesis - reactive oxygen species
12 PPAR signaling pathway
19 Oxidative phosphorylation
8 Steroid biosynthesis
21 Lysosome
31 fatty acid metabolic process
31 lipid transport
20 cellular respiration
20 electron transport chain
17 lipoprotein metabolic process
21 female pregnancy
23 hormone metabolic process
35 lipid localization
19 oxidative phosphorylation
33 alcohol metabolic process
31 lipid catabolic process
25 cellular lipid catabolic process
28 organic acid catabolic process
14 cholesterol biosynthetic process
17 mitochondrial respiratory chain complex assembly
25 steroid biosynthetic process
23 cholesterol metabolic process
35 steroid metabolic process
38 sulfur compound metabolic process
-log10 (P value)

### Figure S5

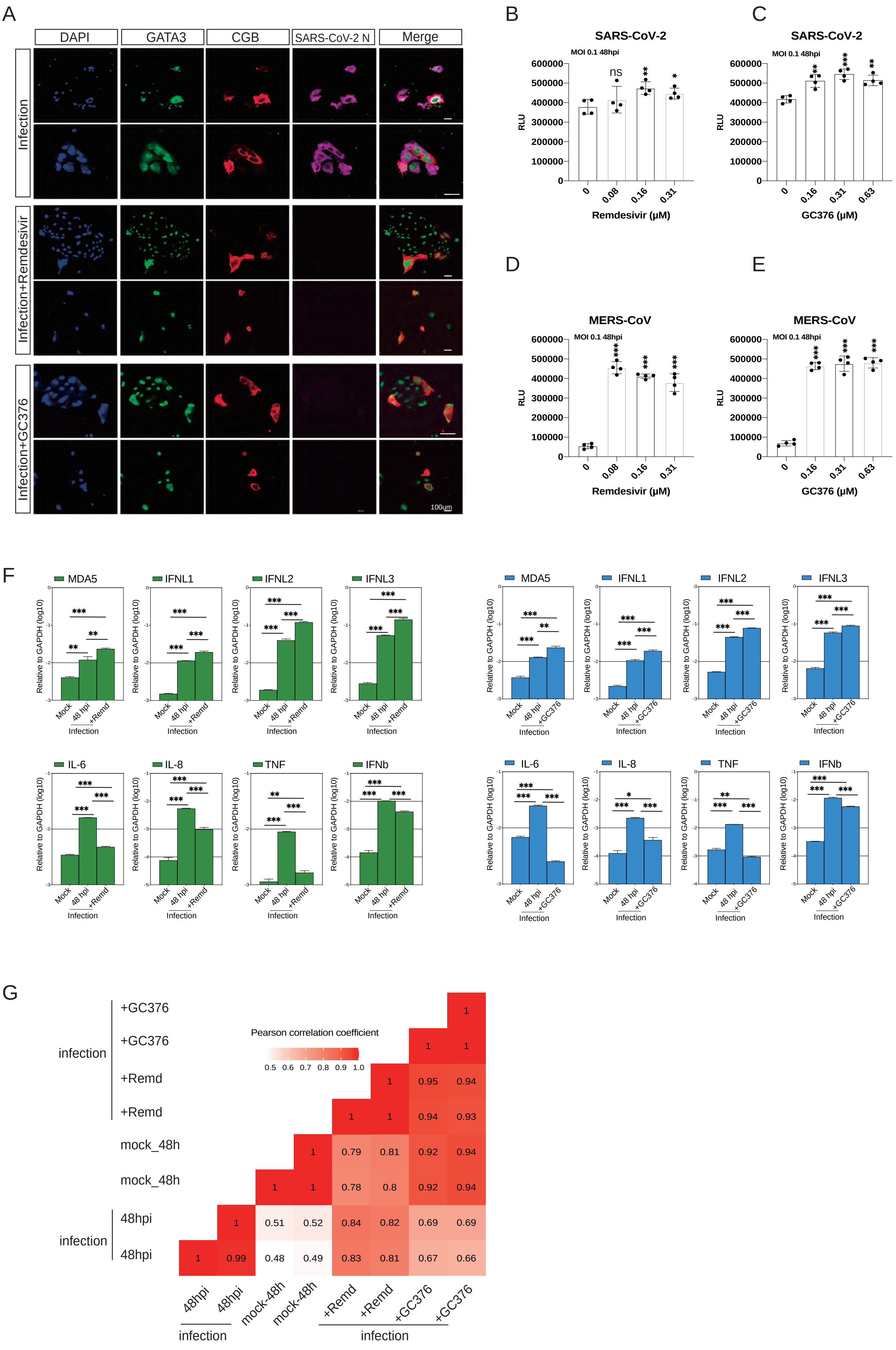

A
DAPI
GATA3
CGB
SARS-CoV-2 N
Merge
Infection
Infection+Remdesivir
Infection+GC376
100um
B
SARS-CoV-2
MOI 0.1 48hpi
RLU
Remdesivir (μM)
C
SARS-CoV-2
MOI 0.1 48hpi
RLU
GC376 (μM)
D
MERS-CoV
MOI 0.1 48hpi
RLU
Remdesivir (μM)
E
MERS-CoV
MOI 0.1 48hpi
RLU
GC376 (μM)
F
MDA5
IFNL1
IFNL2
IFNL3
IL-6
IL-8
TNF
IFNb
Relative to GAPDH (log10)
Mock
48 hpi
+Remd
+GC376
Infection
G
Pearson correlation coefficient
0.5 0.6 0.7 0.8 0.9 1.0
+GC376
+Remd
mock_48h
48hpi
infection

### Figure S6

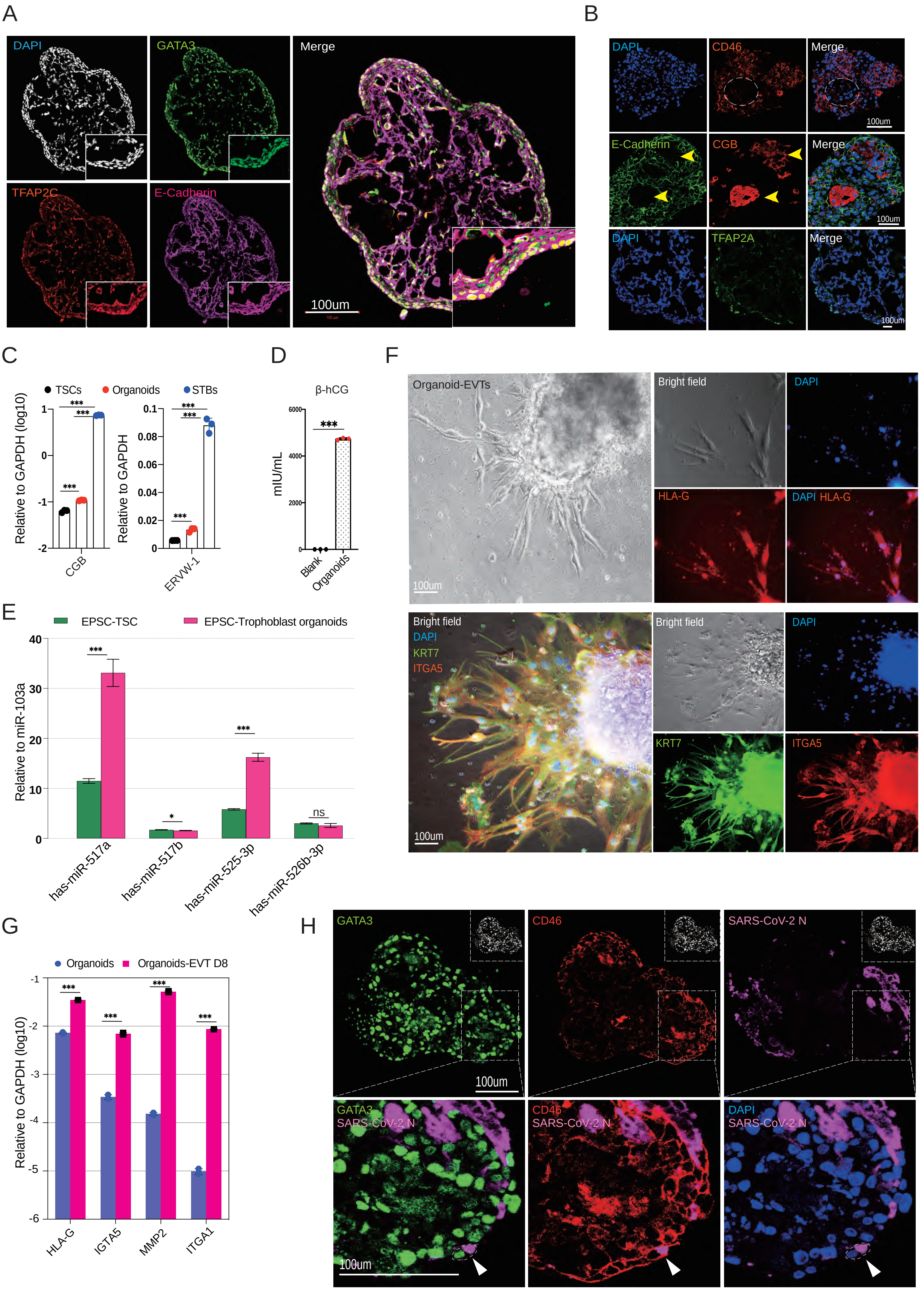

A
DAPI
GATA3
Merge
TFAP2C
E-Cadherin
100um
B
DAPI
CD46
Merge
100um
E-Cadherin
CGB
Merge
100um
DAPI
TFAP2A
Merge
100um
C
TSCs
Organoids
STBs
Relative to GAPDH (log10)
Relative to GAPDH
***
CGB
ERVW-1
D
β-hCG
mIU/mL
Blank
Organoids
F
Organoid-EVTs
Bright field
DAPI
HLA-G
DAPI HLA-G
100um
Bright field
DAPI
KRT7
ITGA5
100um
E
EPSC-TSC
EPSC-Trophoblast organoids
Relative to miR-103a
***
*
ns
has-miR-517a
has-miR-517b
has-miR-525-3p
has-miR-526b-3p
G
Organoids
Organoids-EVT D8
Relative to GAPDH (log10)
HLA-G
IGTA5
MMP2
ITGA1
H
GATA3
CD46
SARS-CoV-2 N
100um
GATA3
SARS-CoV-2 N
CD46
SARS-CoV-2 N
DAPI
SARS-CoV-2 N
100um

### Figure S7

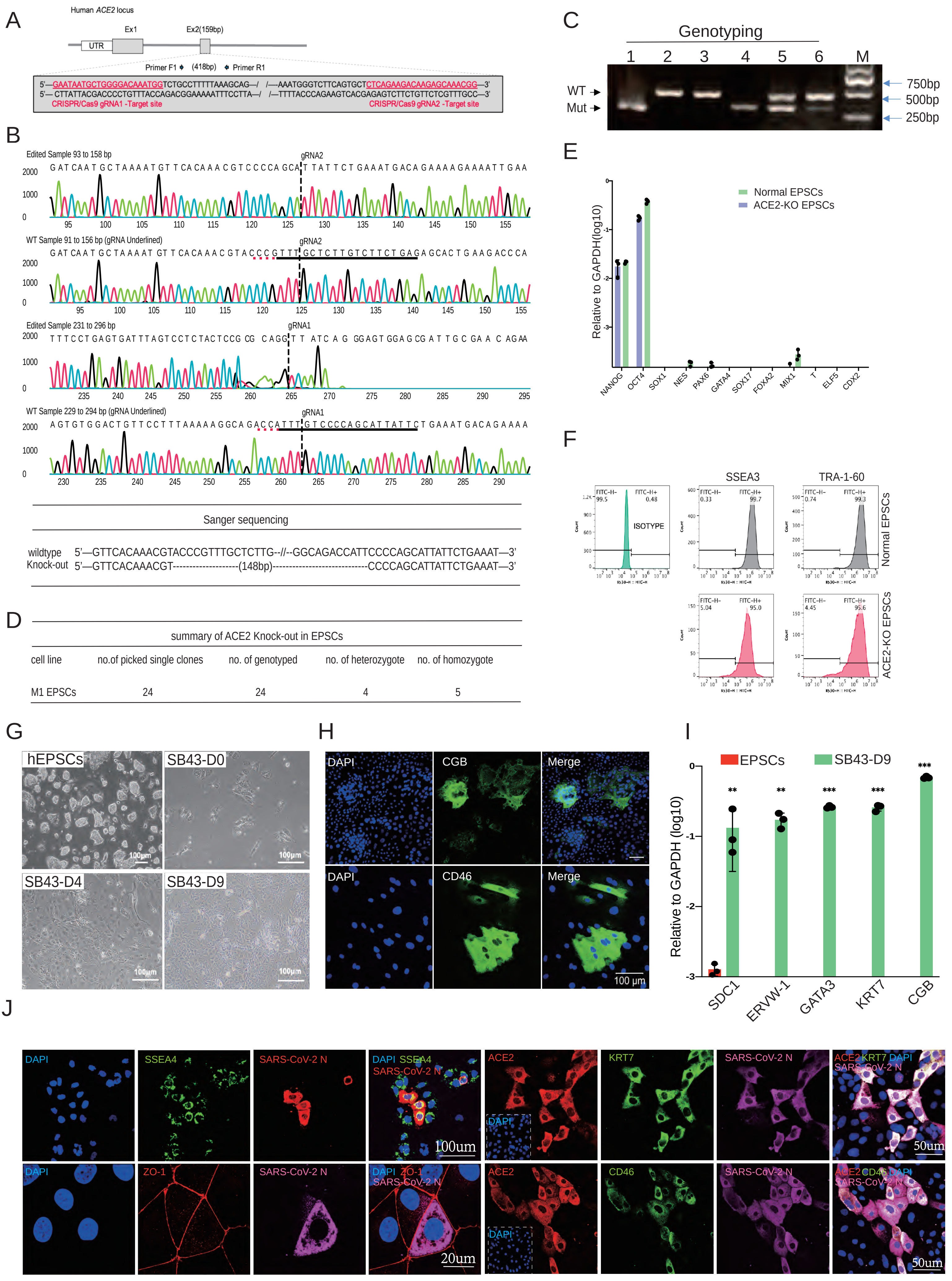
A
Human ACE2 locus
UTR
Ex1
Ex2(159bp)
Primer F1
(418bp)
Primer R1
5'—GAATAATGCTGGGGACAAATGGTCTGCCTTTTTAAAGCAG—/ /—AAATGGGTCTTCAGTGCTCTCAGAAGACAAGAGCAAACGG—3'
5'— CTTATTACGACCCCTGTTTACCAGACGGAAAAATTTCCTTA—/ /—TTTTACCCAGAAGTCACGAGAGTCTTCTGTTCTCGTTTGCC—3'
CRISPR/Cas9 gRNA1 -Target site
CRISPR/Cas9 gRNA2 -Target site
B
Edited Sample 93 to 158 bp
gRNA2
WT Sample 91 to 156 bp (gRNA Underlined)
Edited Sample 231 to 296 bp
gRNA1
WT Sample 229 to 294 bp (gRNA Underlined)
Sanger sequencing
wildtype 5'—GTTCACAAACGTACCCGTTTGCTCTTG--//--GGCAGACCATTCCCCAGCATTATTCTGAAAT—3'
Knock-out 5'—GTTCACAAACGT-------------------(148bp)----------------------------CCCCAGCATTATTCTGAAAT—3'
C
Genotyping
1 2 3 4 5 6 M
WT
Mut
750bp
500bp
250bp
D
summary of ACE2 Knock-out in EPSCs
cell line | no.of picked single clones | no. of genotyped | no. of heterozygote | no. of homozygote
M1 EPSCs | 24 | 24 | 4 | 5
E
Normal EPSCs
ACE2-KO EPSCs
Relative to GAPDH(log10)
NANOG OCT4 SOX1 NES PAX6 GATA4 SOX17 FOXA2 MIX1 T ELF5 CDX2
F
ISOTYPE
SSEA3
TRA-1-60
Normal EPSCs
ACE2-KO EPSCs
G
hEPSCs
SB43-D0
SB43-D4
SB43-D9
100μm
H
DAPI
CGB
Merge
CD46
100 μm
I
EPSCs
SB43-D9
Relative to GAPDH (log10)
SDC1 ERVW-1 GATA3 KRT7 CGB
J
DAPI
SSEA4
SARS-CoV-2 N
100um
ACE2
KRT7
50um
ZO-1
20um
CD46

#### SUPPLEMENTAL FIGURE LEGENDS

**Figure S1. Generating trophoblast stem cells (TSCs) and subtype trophoblasts from hEPSCs**

**A)** RT-qPCR analysis of trophoblast genes ELF5, TFAP2C, TEAD4, TP63 and GATA3 in EPSC-TSCs (passage 10) and TSC-BST (gift from Dr. T. Arima. Passage 3 in Liu lab). Results are normalized to levels of GAPDH using the ΔCt method. Three independent experiments were performed. Data are mean ± SD. ns: not significant (two-tailed unpaired Student's t-test).

**B)** Low HLA-A and -B levels in EPSC-TSCs and TSC-BST in RT-qPCR. Results are normalized to levels of GAPDH using the ΔCt method. Three independent experiments were performed. Data are mean ± SD. *** $p < 0.001$ (two-tailed unpaired Student's t-test).

**C)** Flow cytometry quantification of HLA-A,B,C on EPSC-TSCs and TSC-BST. The antibody isotype is the control.

**D)** RT-qPCR analysis of CDX2 and AME genes MUC16 and GABRP and trophoblast genes ELF5 and GATA3 expression in EPSC-TSC and TSC-BST. Results are normalized to levels of GAPDH using the ΔCt method. Three independent experiments were performed. Data are mean ± SD. ns, not significant (two-tailed unpaired Student's t-test).

**E)** Detection of CDX2 and AME genes in EPSC-TSCs, TSC-BST, and in cells of hEPSCs differentiation toward hTSCs on day 5. AME genes are not detected in established hTSCs. hEPSC differentiation toward TSCs could generate some cells expressing the AME genes. TSC-like colonies from this differentiation were picked and expanded for TSC line establishment.

**F)** Representative bright field images of STBs generated from EPSC-TSCs on day 2, 4 and 6. The arrow points to a multinucleated STB. Scale bar: 100μm

**G)** STBs generated from EPSC-TSCs have distinct morphologies and gene expression. They range from mono-, bi- or multinucleated STBs that express GCM1 and CGB. The multinucleated mature STBs exhibited dotted CGB staining in the cytoplasm. DAPI stains the nucleus. Scale bar: 100μm

**H)** ELISA (IU/L) detection of β-hCG. Supernatant collected from STB cells generated from EPSC-TSCs on Day 2, 3 and 6 are used. Fresh STB medium was used as the blank control.

**I)** Immunofluorescence staining of EVTs for ITGA1 and ITGA5. EVTs are generated from EPSC-TSCs on day 8 differentiation. DAPI stains the nucleus. Scale bar: top panel 100μm; lower panel 50μm

**J)** EVTs generated from EPSC-TSCs (day 8) are immunofluorescence stained for GATA3 and HLA-G. Arrows indicate that a mature EVT (spindle shape) does not express high levels of GATA3. DAPI stains the nucleus. Scale bar: 100μm

**K)** Flow cytometry quantification of EVT marker ITGA5 and ITGA1 on EVTs differentiated from EPSC-TSCs (day 8).

**L)** Detection of cell type specific genes in RNAseq analysis of EPSC-TSCs, and STBs and EVTs generated from EPSC-TSCs. z-score normalized cpm are used. Genes are ordered by z-scores.

**M)** Expression of HLA genes in the recently published human placenta-derived trophoblasts (Sheridan, M.A., 2021) and in EPSC-TSCs and their derivative trophoblasts. Error bars: standard errors.

**Figure S2. Trophoblasts derived from hEPSCs resemble those in the human embryos and the placenta**

**A)** Violin plots of trophoblast gene expression TEAD4, TP63, CGB5, CSH2, HLA-G and MMP2 (log-transformed TPM) in human peri-implantation embryo and placenta trophoblasts as shown in Figure 2B.

**B)** Violin plots of expression of AME genes MUC16, GABRP, VTCN1, ITGB6, ISL1 and CDX2 (log-transformed TPM) in human peri-implantation embryo and placenta trophoblasts.

**C)** Heatmap of Pearson correlation coefficients of peri-implantation embryo and placenta trophoblast pseudobulk transcriptomics and the *in vitro* cultured trophoblasts bulk RNAseq, based on regressed whole-transcriptomic expression. Pseudobulk samples are assembled by 50 single cells from each peri-implantation embryo and placenta trophoblast clusters. *In vitro* trophoblast samples include blastocyst- and placenta-derived TSCs (TSC-BST, TSC-ct) and the trophoblast subtypes STBs (STB-BST, STB-ct) and EVTs (EVT-BST, EVT-ct), and EPSC-TSCs and the trophoblast subtypes STBs and EVTs. Each row or column represents one sample's correlation coefficients with all other samples.

**D)** Upper panel: Scatter plots showing the positive Pearson correlations of CD46, CGB5, ENG and CSH2 with ACE2 expression. Linear regression line is drawn in black dashed line. Cell density is represented by the darkness of blue. Lower panel: Scaled expression of STB genes CD46, CGB5, ENG and CSH2 in human peri-implantation embryo and placenta trophoblasts.

**E)** Scaled expression of TMPRSS2, BSG and AXL in human peri-implantation embryo and placenta trophoblasts.

**F-G)** RNA-seq signal of ACE2 in trophoblasts. *In vivo* origin: TSC-BST, TSC-ct, and their differentiated trophoblast subtypes (STB/EVT-BST/ct). The ACE2 locus genomic region is plotted at the bottom, where each vertical bar represents an exon, and the transcription direction goes from right to left.

**Figure S3. eSTBs are highly susceptible to SARS-CoV-2 infection**

**A)** RT-qPCR detection of SARS-CoV-2 genome copy number in cell lysates of 2 hpi and 48 hpi EPSCs and EPSC-TSCs. Data are mean $\pm$ SD. ns, not significant, *** $p < 0.001$ (two-tailed unpaired Student's t-test between 2 hpi and 48 hpi).

**B)** Immunofluorescence staining of 48 hpi hEPSCs for pluripotency markers and SARS-CoV-2 N protein. hEPSCs do not express ACE2 and are not susceptible to the infection. Scale bar: 100μm.

**C)** Immunofluorescence staining of 48 hpi EPSC-TSCs for SARS-CoV-2 N protein. Scale bar: 100 mm. Right panel: Percentages of SARS-CoV-2 N protein positive EPSC-TSCs. Error bar: mean and standard error (SEM). n=36, quantification of 36 random immunofluorescence staining images.

**D)** RT-qPCR analysis of SARS-CoV-2 genome copy number in supernatants and cell lysates of 2, 24, 48 and 72 hpi EVTs. Cells were submitted for SARS-CoV-2 co-incubation on day 8 of EPSC-TSC differentiation toward EVTs. Data are mean + SD. *** $p < 0.001$ (two-tailed unpaired Student's t-test).

**E)** Immunofluorescence staining of 48 hpi EVTs for SARS-CoV-2 N protein. Scale bar: 100 mm. Right panel: Percentages of SARS-CoV-2 N protein positive EVTs. Error bar: mean and standard error (SEM). n=36, quantification of 36 random immunofluorescence staining images.

**F)** RT-qPCR analysis of SARS-CoV-2 genome copy number in cell lysates of 48 hpi EPSC-TSCs and STB-D4 by RT-qPCR. Gene expression levels were normalized to GAPDH using the ΔCt method. Datapoints are mean and SD from three independent experiments. *** $p < 0.001$ (two-tailed unpaired Student's t-test).

**G)** Immunofluorescence staining 48 hpi STB-D6 for ACE2, CGB and SARS-CoV-2 N protein, and for early STB markers CD46 and SSEA4. Top panel: ACE2 is not or lowly expressed in the $CGB^+$ multinucleated mature STBs, which are rarely infected by SARS-CoV-2. The infected cells are usually positive for ACE2, CD46 and SSEA4. Lower panels: arrowheads indicate that some CGB-low, mono-or bi-nucleated cells could be infected by SARS-CoV-2. DAPI stains the nucleus. Scale bar: 50μm and 100μm as specified in the images.

**H)** Percentages of mono-, bi- and multi-nucleated $CGB^+$ cells in SARS-CoV-2 infected STB-D6.

**I)** Quantification of percentages of mono-, bi- and multi-nucleated cells in STB-D2. Error bar, mean and standard error (SEM). n=40, quantification of 40 random immunofluorescence staining images, total 6734 cells.

**J)** Immunofluorescence staining for CGB in STB-D2. Dotted box showed a multinucleated $CGB^+$ cells. Scale bar: 100 μm. Right panel: Quantification of the percentages of $CGB^+$ in STB-D2. Error bar, mean and standard error (SEM). n=8, quantification of 8 random immunofluorescence staining images, total 5820 cells.

**K-L)** RNAseq analysis of expression of STB genes CD46 and CGB5 and of ACE2 and TMPRSS2 in EPSC-TSCs and EPSC-TSC differentiation toward STBs at different time points.

**M)** RNAseq signal of ACE2 gene in EPSC-TSCs and in their differentiation toward STBs at different time points. The ACE2 genomic region is plotted at the bottom where each vertical bar represents an exon. The transcription direction goes from right to left.

**Figure S4. SARS-CoV-2 infection impairs differentiation toward STBs**

**A)** Heatmap for correlation between infected STBs and mock infection STBs. Unexpressed genes are filtered out and cpm is used to calculate Pearson correlation coefficient.

**B)** Venn plots for upregulated genes and downregulated genes after virus infection in STB-D3 and STB-D4.

**C)** The top 50 differentially expressed genes in STB-D3 and STB-D4 after virus infection. Z-score of log2 transformed cpm was used.

**D)** Gene set enrichment analysis (GSEA) for apoptosis pathway in STB-D3 and STB-D4 after virus infection. Red, upregulated genes; blue, downregulated genes.

**E)** RNAseq signal of HERV-K and HERV-FRD in infected STB-D3 and STB-D4 and the mock controls. Library size is used to normalize the reads.

**F)** Validation of the machine learning model (linear regression) for prediction of pseudotime. Drawn dots are the 895 *in vivo* cells from PI-TB to PI-STB subtypes (25% of all cells in the two subtypes). The other 75% cells are used to train the linear regression model. X-axis is the predicted pseudotime value by the linear regression model, and y-axis is the “observed pseudotime” computed by using SCANPY. Darkness of the blue color indicates the density computed by Gaussian kernel. Red dots are the outliers. A coefficient of 0.976 indicates high alignment of predicted values and observed values, and that the trained model could be further used to predict the pseudotime for *in vitro* cells.

**G)** GO and KEGG analysis for shared upregulated and downregulated genes in infected STB-D3 and STB-D4.

**Figure S5. Low dosages of antiviral drugs effectively eliminated SARS-CoV-2 and MERS-CoV infection of eSTBs**

**A)** Immunofluorescence staining for GATA3, CGB and SARS-CoV-2 N protein in 48 hpi eSTBs treated with remdesivir or GC376. N protein is not detectable in the drug-treated cells. Scale bar: 100μm

**B-C)** CPE (cytopathogenic effects) inhibition assay of SARS-CoV-2 48 hpi eSTBs after different concentrations of remdesivir (B) or GC376 (C) treatment. Shown is the luminance reading by CellTiter-Glo. (Values are mean + SEM from 3 separate experiments. ns, not significant; *p < 0.05; **p < 0.01; *** p < 0.001 (two-tailed unpaired Student's t-test).

**D-E)** CPE inhibition assay of MERS-CoV 48 hpi eSTBs after different concentrations of remdesivir (D) or GC376 (E) treatment. Shown is the luminance reading by CellTiter-Glo kit measured at 48hpi. (Values are mean ± SEM from 3 separate experiments, *** p < 0.001 (two-tailed unpaired Student's t-test).

**F)** RT-qPCR detection of gene expression changes of MDA5, IFNLs, IL6, IL8, TNF and IFNb in SARS-CoV-2 infected eSTBs, in the presence or absence of remdesivir or GC376. (Data are mean ± SD, n=3 independent replicates from 3 separate sample extracts. *p < 0.05; ** p < 0.01; *** p < 0.001 (two-tailed unpaired Student's t-test). Gene expression levels were normalized to GAPDH using the ΔCt method.

**G)** The heatmap for correlation between infected and mock infection eSTBs. Non-expressed genes were filtered out and cpm was used to calculate the Pearson correlation coefficient.

**Figure S6. Infection of EPSC-TSC trophoblast organoids by SARS-CoV-2**

**A)** Immunofluorescence staining of trophoblast organoid cryosections for TFAP2C, GATA3 and E-cadherin. Scale bar: 100μm

**B)** Immunofluorescence staining of trophoblast organoid paraffin sections for CD46, E-cadherin, TFAP2A and CGB. Dash line circles indicate multinucleated STBs, which are also indicated by yellow arrowheads in the E-Cadherin and CGB staining panels. Scale bar: 100μm

**C)** Expression of STB genes (ERVW-1 and CGB) in STBs, EPSC-TSCs and trophoblast organoids by RT-qPCR (Data are mean $\pm$ SD, n=3 independent replicates from 3 separate sample extracts. *** $p < 0.001$ (two-tailed unpaired Student's t-test). Gene expression was normalized to GAPDH using the ΔCt method.

**D)** ELISA (mIU/mL) detection of hCG-β secreted by trophoblast organoids in the supernatant. Fresh trophoblast organoid medium was used as the blank control. n=3 independent replicates. *** $p < 0.001$ (two-tailed unpaired Student's t-test).

**E)** Trophoblast-specific C19MC miRNAs (has-miR-517a, 517b, 525-3p and 526-3p) expression levels (RT-qPCR) in EPSC-TSCs and EPSC-TSC derived trophoblast organoids. miRNA103a expression is used as the control. (Data are mean $\pm$ SD, n=3 independent replicates from 3 separate sample extracts. ns, not significant, * $p < 0.05$, *** $p < 0.001$ (two-tailed unpaired Student's t-test).

**F)** Representative phase-contrast image of cells migrating out of a trophoblast organoid seeded in EVT medium for 8 days. EVTs were stained for HLA-G, ITGA5 and KRT7. Nuclei were counterstained with DAPI. Scale bar: 100μm

**G)** Expression of EVT genes HLA-G, ITGA5, MMP2 and ITGA1 in trophoblast organoids and trophoblast organoid-derived EVTs analyzed by RT-qPCR. (Data are mean $\pm$ SD, n=3 independent replicates from 3 separate sample extracts. *** $p < 0.001$ (two-tailed unpaired Student's t-test). Gene expression was normalized to GAPDH using the ΔCt method.

**H)** Additional immunofluorescence staining images of SARS-CoV-2-infected trophoblast organoid paraffin sections for GATA3 and SARS-CoV-2-N. The infected cells were primarily found at the periphery of the organoids where trophoblast progenitors or eSTBs were located. Higher resolution images in the lower panel are from the dash line box area. Nuclei were counterstained with DAPI. Arrows indicate an infected cell that is co-stained for GATA3, CD46 and SARS-CoV-2-N. Scale bar: 100μm

**Figure S7. ACE2 is required for SARS-CoV-2 infection of trophoblasts differentiated from hEPSCs**

**A)** CRISPR/Cas9-mediated knockout of the human ACE2 gene. A pair of gRNAs targeting exon 2 of ACE2 are listed in red.

**B)** Sanger sequencing of the mutant PCR fragments reveals a 148bp deletion between the two CRISPR gRNAs in exon 2 as expected.

**C)** Genotyping of ACE2-KO hEPSC colonies by genomic DNA PCR. Colonies 1 and 4 are homozygous KO mutants.

**D)** Genome editing efficiency at the ACE2 locus in hEPSCs. Five out 24 genotyped colonies were the homozygous mutant ones.

**E)** Expression of pluripotency and various cell lineage genes in normal and ACE2-KO hEPSCs.

**F)** Flow cytometry quantification of pluripotency markers SSEA3 and TRA-1-60 on normal and ACE2-KO hEPSCs.

**G)** Morphology of hEPSCs and SB43-treated cells on days 0, 4 and 9. Scale bars: 100μm

**H)** Representative immunofluorescence staining of STB markers CGB and CD46 in SB43-treated hEPSCs on day 9. Scale bars: 100μm

**I)** Expression of trophoblast genes in SB43-treated hEPSCs on day 9 by RT-qPCR. (Data are mean $\pm$ SD, n=3 independent replicates from 3 separate sample extracts. ** $p < 0.01$, *** $p < 0.001$ (two-tailed unpaired Student's t-test). Gene expression was normalized to GAPDH using the $\Delta$Ct method.

**J)** Representative immunofluorescence images of SARS-CoV-2 infection in SB43-treated hEPSCs (day 4). KRT7 is a pan-trophoblast marker. CD46 and SSEA4 stain early STBs. ZO-1 stains cell tight junction. SARS-CoV-2 N protein was detected by immune serum. More mononucleated cells were infected. Scale bar: 50μm and 100μm as specified in the images.

#### STAR * METHODS

#### RESOURCE AVAILABILITY

##### Lead contact

##### Materials availability

All stable reagents generated in this study are available from the lead contact.

##### Data and code availability

RNA-seq data generated in this study have been deposited at NCBI Gene Expression Omnibus (GEO): GSE190432.

#### EXPERIMENTAL MODEL AND SUBJECT DETAILS

##### Human expanded potential stem cells (hEPSCs) and cell lines

Human embryonic stem cells (hESCs) Man-1/M1 were converted to hEPSCs, as was previously described[1]. Human EPSC cultures were routinely maintained on STO feeders. Irradiation inactivated STO cells were prepared 3-4 days before seeding hEPSCs on 0.1% gelatinised plates at the density of ~$3.125\times10^4$ cells/cm2. STO cells were maintained in regular M10 medium: knockout DMEM, 10% FBS, 1×Glutamine Penicillin-Streptomycin and 1× Minimum Essential Medium (MEM) Vitamin Solution. Human colon Caco-2 cells and monkey Vero E6 cells were maintained in DMEM culture medium supplemented with 10% heat-inactivated FBS, 50 Uml−1 penicillin and 50 μgml−1 streptomycin. All cells were maintained in a 5% $CO_2$ incubator at 37 ℃ and routinely tested for mycoplasma.

##### Virus

The SARS-CoV-2 HKU-001a strain (GenBank accession number: MT230904) was isolated from the nasopharyngeal aspirate specimen of a patient who was laboratory-confirmed to have COVID-19 in Hong Kong[2]. The MERS-CoV strain (HCoV-EMC/2012) was a gift from R. Fouchier. All experiments involving live SARS-CoV-2 and MERS-CoV followed the approved standard operating procedures of the biosafety level 3 facility at the University of Hong Kong.

#### METHODS DETAILS

##### Culture of hEPSCs

Human EPSC cells were maintained on STO feeder layers and enzymatically passaged (1:10) every 3-5 days by a brief PBS wash followed by treatment with TrypLE (Gibco. Cat. 12605036) for 5 minutes. Cells were dissociated and centrifuged (300g for 3 min) in 10% foetal bovine serum (FBS)-containing medium (M10 medium). After removing supernatant, human EPSCs were resuspended and seeded in hEPSC Medium (EPSCM) supplemented with 5.0 µM Y27632 (Tocris. Cat. 1254). hEPSCM are N2B27-based media supplement with small molecular as previously published[1]. N2B27 basal media [1:1 of DMEM/F12 (Thermo, Cat.21331020) and Neurobasal Medium (Thermo, Cat. 21103049); 200x N2 supplement and 100x B27 supplement].

##### Differentiation of hEPSCs to trophoblast lineages by the TGF-β inhibitor SB431542

Human EPSCs were dissociated with TrypLE (Gibco. Cat. 12605036) and seeded in 100x Geltrex (Thermo. Cat. A1413302) coated six-well plates at a density of $1\times10^5$ cells per well. Cells were cultured (pre-treatment) in 20% KSR media supplemented with 10µM Y27632 for one day. From the second day, 10µM SB431542 (Tocris. Cat. 1614) was added into 20% KSR media to start the differentiation. Cells were collected at the indicated time points for analysis.

**Derivation of human Trophoblast Stem Cell (hTSCs) from hEPSCs**

Single cell-dissociated hEPSCs were plated on 6-well plates pre-coated with 100x Geltrex (Thermo. Cat. A1413302) at a density of 2,000 cells per well and cultured in hTSC media: DMEM/F12 (Gibco. Cat. 21331-020) supplemented with 50.0uM β-mercaptoethanol (Thermo. Cat. 31350010), 0.2% FBS (Gibco. Cat. 10270), 0.5% Penicillin-Streptomycin-Glutamine (Thermo, Cat.10378016), 0.3% BSA (Gibco. Cat. 15260037),1.0% ITS-X supplement (Gibco. Cat. 51500056), 50.0 µg/mL Vc (Sigma. Cat. 49752-100G), 50.0 ng/mL EGF (Thermo Fisher, PHG0311), 2.0 µM CHIR99021 (GSK3i. Tocris. Cat. 4423), 0.5 µM A83-01 (Tocris. Cat. 2939), 1.0 µM SB431542 (Tocris. Cat. 1614), 10.0 µM VPA (Stemcell. Cat. 72292) and 5.0 µM Y27632 (Tocris. Cat. 1254). After 12-14 days of culture, the colonies with TSC-like morphologies were picked, dissociated in TrypLE (Gibco. Cat. 12605036) and replated on a plate pre-coated with 100x Matrigel. After 4-5 passages, the cells were collected for syncytiotrophoblast (STB) and extravillous trophoblast (EVT) differentiation[3].

**Differentiation of EPSC-TSCs to STBs**

Wells of a six-well plate were coated with 100× Matrigel (Corning. Cat. 354230) for at least 1h. $1.0\times10^5$ hTSCs were seeded per well in 2 mL STB medium [DMEM/F12 (Thermo, Cat.21331020) supplemented with 50uM β-mercaptoethanol, 0.5% Penicillin-Streptomycin-Glutamine (Thermo, Cat.10378016), 0.3% BSA,1% ITS-X, 2.5 µM Y-27632, 2µM Forskolin (Sigma-Aldrich. Cat. F3917), and 4% KnockOut Serum Replacement (Thermo. Cat. 10828028)]. Media was changed on day 3, and the cells were ready for downstream analysis on day6.

**Differentiation of EPSC-TSCs to EVTs**

For EVT differentiation, wells of a 6-well plate were coated with 100× Matrigel (Corning. Cat. 354230) for at least 1h. 1.0 ×105 hTSCs were seeded per well in 3.0 mL EVT basal medium [DMEM/F12 (Thermo, Cat.21331020) supplemented with 50uM β-mercaptoethanol, Penicillin-Streptomycin-Glutamine (Thermo, Cat.10378016), 0.3% BSA, 1%ITS-X, 7.5uM A83-01, 2.5uM Y27632] supplemented with 4%,100 ng/mL NRG1 (Cell signalling. Cat. 5218SC) and 2% Matrigel. On day 3, the media were replaced with 2 mL EVT basal medium supplemented with 4% KSR and 0.5% Matrigel. On day 6, the media were replaced with 2 mL EVT basal medium, and Matrigel was added to a 0.5% final concentration. On day 8, the cells were ready for downstream analysis.

**Generation of trophoblast organoids from EPSC-TSCs**

EPSC-TSCs were digested with TrypLE (Gibco. Cat. 12605036) and dissociated to single cells by pipetting. After centrifugation, growth factor-reduced Matrigel (GFR-M. Corning) was added to reach a final concentration of 60% and the rest 40% was trophoblast organoid medium (TOM). Matrigel/TOM (40 uL) containing $1.0 \times 10^5$ hTSCs was seeded into the well centre of a 24-well plates. After 3 mins at 37℃ in a $CO_2$ incubator, the plates were turned upside down to ensure equal spreading of the cells in the well. After 15 min at 37℃ in an incubator, the solidifying GFR-M formed domes and were carefully overlaid with 500 uL TOM. Trophoblast organoids were allowed to form for 4–6 days at P0. After 20 days culturing, trophoblast organoids were collected by dissolving Matrigel with recovery solution (Corning. Cat. 354253) and submitted to SARS-CoV-2 infection. TOM is composed of N2B27 basal media, recombinant human EGF 50 ng/mL, CHIR99021 1.5µM, recombinant human R-spondin-1 80 ng/mL, recombinant human FGF-2 100 ng/mL, recombinant human HGF 50 ng/mL, A83-01 500 nM, Prostaglandin E2 2.5 µM, Y-27632 2 µM. Store the medium at 4 °C for up to 2 weeks.

**Other human cell lines for SARS-CoV-2 infection**

Human colon epithelial (Caco-2) cells and African green monkey kidney (Vero E6) cells were purchased from ATCC. All cells cultured in this study were maintained at 37℃ with 5% $CO_2$, unless stated otherwise, and had been confirmed to be free of mycoplasma contamination by PlasmoTest (InvivoGen).

**SARS-CoV-2 infection and detection**

The SARS-CoV-2 strain (SARS-CoV-2 HKU-001a; GenBank accession number MT230940) was isolated from a nasopharyngeal aspirate specimen from a COVID-19 patient in Hong Kong. SARS-CoV-2 stock was propagated using Vero E6 cells, and the titer of supernatant was assessed by plaque assays. All experiments involving with live SARS-CoV-2 followed the approved standard operating procedures of Biosafety Level 3 facilities in Queen Mary Hospital, The University of Hong Kong.

Indicated hEPSCs were seeded one day before infection and other types of differentiated cells at the suitable time point. On the day of infection, cells were washed with PBS and infected at the indicated multiplicity of infection (MOI) by diluting viruses in basal medium. Cells were incubated at 37°C for 2 hrs. Subsequently, the inoculum was removed, replaced with complete culture medium, and further incubated until harvest. Cytopathic effects (CPE) were monitored daily by light microscopy, and cell supernatant and lysates at indicated time point were collected for RT-qPCR to assess the viral RNA load, which had been calibrated to viral load determination by plaque assays.

**Detection of SARS-CoV-2 virus**

For viral detection, the supernatants of the cultured cells challenged by SARS-CoV-2 were harvested at various time points. A total of 140 μL of culture supernatant was lysed with 560 μL of AVL buffer, which was subsequently extracted for total RNA with the QIAamp® Viral RNA Mini Kit (QIAGEN. Cat. 52906).

For virus replication kinetics assays, the extracted RNA was quantified with the one-step QuantiNova Probe RT-PCR kit (QIAGEN. Cat. 208354). Each 20 μL reaction mixture contained 10 μL of 2× QuantiNova Probe RT-PCR Master Mix, 0.2 μL of QuantiNova Probe RT-Mix, 1.6 μL each of 10μM forward and reverse primer, 0.4 μL of 10 μM probe, 5 μL of extracted RNA as template, and 1.2 μL of RNase-free water. Reactions were incubated at 45 °C for 10 min for reverse transcription, 95 °C for 5 min for denaturation, 45 cycles of 95 °C for 5 s and 55 °C for 30 s, followed by a cooling step at 40 °C for 30 s. The primers and probe sequences were against the RNA-dependent RNA polymerase/ helicase (RdRP/Hel) gene region of SARS-CoV-2 were listed in Supplementary Table S3.

**Plaque assay**

Plaque assay was performed as we previously described (Yuan et al, 2021)[4]. Briefly, Vero E6 cells were seeded at 300,000 cells/well in 12-well tissue culture plates on the day before carrying out the assay. After 24 h of incubation, a serial dilution of supernatant were added to the cell monolayer and the plates were further incubated for 1 h at 37 C in 5% $CO_2$ before removal of unbound viral particles by aspiration of the media and washing once with DMEM. Monolayers were then overlaid with media containing 1% low melting agarose (Cambrex Corporation, East Rutherford, NJ, USA) in DMEM, inverted and incubated as above for another 72 h. The wells were then fixed with 10% formaldehyde (BDH, Merck, Darmstadt, Germany) overnight. After removal of the agarose plugs, the monolayers were stained with 0.7% crystal violet (BDH, Merck) and the plaques counted. The plaque assay experiments were performed in triplicate.

**Guide RNA design, RNA synthesis and plasmid DNA preparation**

The human ACE2 exon 2 sequence was analysed using the online CRISPOR tool for designing a pair of highly-specific gRNAs. Chemically-synthesized ssDNA oligos were incubated at 95°C for 10 mins for annealing into dsDNAs, which were then ligated into a linearized empty gRNA vector using the DNA Ligation Kit, Mighty Mix (TaKaRa. Cat. 6023). The ligation product was transformed into Chemically Competent DH 5α Cell (KT Health) and spread onto a LB agar plate with ampicillin for selection. The next day, single colonies were picked and cultured in LB broth with ampicillin for plasmid miniprep using the TIANprep Rapid Mini Plasmid Kit (Tiangen. Cat. DP105). After confirmation of ligated gRNA sequences by Sanger Sequencing, gRNA plasmids with the correct target sequence was amplified and extracted by the Endofree Plasmid Kit II (Tiangen. Cat. DP118) to purify a large amount of endotoxin-free gRNA plasmid DNA for electroporation. Plasmid DNA of BSD-Cas9 was similarly prepared as described previously[1].

**Electroporation and selection**

hEPSCs were cultured on hBN feeder. Electroporation was performed when the cells reached 70%-80% confluence. Plasmid DNA including Cas9 and double sgRNA with ration 2:1:1 (4ug Cas9, 2ug sgRNA1, 2ug sgRNA2 per 1millon cells as one group) were added into opti-MEM (Gibico. Cat. 31985062). hEPSCs were washed twice with PBS, then dissociated into single cells using 0.05% Trypsin-EDTA.

M10 medium (DMEM with 10% FSB) was added to neutralize trypsin. Single cell hEPSCs were re-suspended in Opti-MEM medium containing DNA mixture. Electroporation was performed using Bio-Rad Gene Pulser Xcell Electroporation Systems using 0.4cm cuvette with 230V, 500uF. After electroporation, cells were seeded with recovery medium (500 ul EPSCM + 10% KSR+10uM Y27632). After incubation overnight, cells were switched to normal EPSC medium. One day after electroporation, cells were selected by 10ug/mL Blasticidin S HCl (Thermo. Cat. A1113903). Two days after electroporation, cells were selected by 10ug/mL Blasticidin S HCl and 1ug/ml Puromycin Dihydrochloride (Thermo. Cat. A1113802) for another 2 days. After 7-10 days, single colonies were picked, expanded, and genotyped.

**Genotyping and Sanger sequencing**

Single colonies were picked and digested into single cells in a 96-well plate by 0.05% trypsin-EDTA for 3-5 mins. Half of these cells were transferred into a 48 well plate with EPSCM medium and SNL feeders for culturing. The other half cells of the same colony were for genotyping which was performed by using primers designed to amplify the targeted band as well as the wild types. The genotyping primers according to targeted ACE2 were as following: Forward primer: 5'-GTGGCCTGGTCACTCTTAAC-3'; Reverse primer: 5'-CAAATAAAGGCAGCTGCTGTG-3'. The mutant PCR band was gel-purified and confirmed by Sanger sequencing for 148bp deletion.

**Reverse Transcription- Quantitative Polymerase Chain Reaction (RT-qPCR)**

Total RNA extraction was performed using the RNeasy Mini Kit (Qiagen. Cat. 74104) as per manufacturer's instructions. The isolated RNA was reverse transcribed into complementary (cDNA) using the Fastking gDNA Dispelling RT SuperMix (Tiangen. Cat. KR118) on a thermal cycler. The PowerUp™ SYBR™ Green Master Mix (Applied Biosystems) was used for checking intracellular SARS-CoV-2 and gene expression. Primer sequences are listed in Supplementary Table S3. All qPCR experiments were performed using The StepOnePlus™ Real-Time PCR System (Applied Biosystems). Viral RNA load and gene expression levels were normalized to GAPDH using the ΔCt method. Data were analysed using one/two-tailed Student's t-test on Prism 8 (GraphPad).

**Immunofluorescence staining**

The samples were fixed in 4% paraformaldehyde (Sigma. Cat. P6148) at room temperature for 15 minutes, permeabilized with 0.3% Triton-X100 (Sigma. Cat. T8787) for 30 minutes blocked for 3 hrs with 10% donkey serum (Sigma. Cat. D9663) and 1% BSA (Sigma. Cat. A2153) and followed by an incubation with primary antibodies for overnight in a 4℃ cold room. Fluorescence-conjugated secondary antibodies were used to incubate the samples at room temperature for 1 hr. After antibody staining, samples were counter-stained with 10 µg/ml DAPI (Thermo Fisher Scientific. Cat. 62248) for 10 mins to mark nuclei and were observed under a fluorescence microscope.

**Western blotting**

Proteins were separated on 7.5% polyacrylamide gels (Bio-Rad. Cat. 1610180) and transferred to PVDF membrane in Bio-Rad transblot turbo system according to manufacturer's guidance. The following primary antibodies were used: rabbit ACE2 (1:500, Abclonal. Cat. A4612), rabbit β-actin (1:5000, Abmart. Cat. P30002M). Goat anti-rabbit IgG H&L (HRP) (1:10000, Abcam. Cat. ab205718) was used as the secondary antibody. The image was developed and analyzed by ChemiDoc Imaging System.

**Flow cytometry**

hEPSCs were digested with 0.25% trypsin/EDTA for 2-3 minutes at 37 °C and dissociated to single cells by pipetting. The dissociated cells were filtered with a 40 µm nylon mesh (Corning. Cat. 352235) to remove clumps. After centrifugation, the cells were fixed using Fixation Medium according to the manufacturers' manual (BD Cytofix. Cat.554655) and the washed cells were stored at 4 °C in PBS supplemented with 0.1% NaN3 (Sigma. Cat. 199931) and 5% FBS (Gibco. Cat. 10270) before analysed with flow cytometry. All the samples were analysed using ACEA NovoCyte Quanteon. 488nm (530/30 bandpass filter) and 561nm (610/20 bandpass filter) channels were used to detect FITC and excluded autofluorescence. 405nm (445/45 bandpass filter) channel was used to detect DAPI positive cells. FACS data were analysed by Flowjo software.

**Transwell invasion assay**

The invasion ability of TSC derived EVTs were determined by cell invasion (354480, Corning, city, USA) according to manufacturers' instructions. Briefly, the invasion chambers were incubated with warm DMEM base medium at 37℃ for 1 h. After rehydration, the medium was removed. ETSC-EVTs (1x10$^5$ cells/well) was prepared in DMEM basal medium. The mixture was added into an invasion chamber and placed into a 24 well culture plate, with the lower chamber filled with DMEM containing 10% FBS. The cells were allowed to pass through the chamber and attached to the lower bottom of the polycarbonate membrane for 22 h. After that, the medium from the top insert was aspirated, and non-invasive/-migratory cells on the upper surface were wiped away with a cotton bud. The invaded/migrated cells on the lower surface were stained with crystal violet for 15 min. The membrane was observed under a light microscope.

**Statistical analysis and reproducibility**

The statistical analysis was conducted with Microsoft Excel or Prism 8 (GraphPad). P values were calculated using (un)paired Student's t-test. The figure legends have the exact number of measurements, the number of independent experiments and the statistical test used for each analysis performed. Experiments were repeated independently with similar results obtained.

**RNA-seq analysis**

Cutadapt was used to remove adapter sequences and low-quality 3′ end sequences. Processed reads were mapped to the human hg38 genome assembly by hisat2. Gene annotation form Ensembl was used. FeatureCounts was used to quantify gene expression. Genes with mean count number <5 were filtered out. Transposable element annotations were from UCSC Genome Browser (RepeatMasker). SQuIRE with "total" mode was used to quantify TE expression. DESeq2 was used to analysed differentially expressed genes and TEs. Genes and TEs with expression fold change > 1.5 and adjusted P-value < 0.05 were considered significant differential expressed. R package clusterprofiler was used for gene ontology (GO) and KEGG analysis. GSEA software (Gene Set Enrichment Analysis) was performed by GSEApy and the gene sets used were downloaded directly from https://www.gsea-msigdb.org. Bigwig files for RNA-seq signal were generated by bamCoverage from Deeptools and IGV was used for visualization. For the RNA-seq signal on endogenous retrovirus, GenBank accession number AY101582.1, BC068585.1 and N675077.1 were used to inquire the sequence of HERV-W, HERV-FRD and HERV-K and the reads were mapped by hisat2. GenBank accession number MN985325 was used to inquire the sequence of SARS-CoV-2 genes, then the reads were mapped to sequences by hisat2 and quantified. For data deposited in E-MTAB-10429, processed count table was used directly.

**scRNA Preprocessing**

scRNA preprocessing was performed according to the SCANPY pipelines[5]. Briefly, Transcripts Per Million (TPM) of pre- to post- implantation cells were extracted from Zhou's dataset[6] (GEO accession number: GSE109555), and placenta cells were extracted from Liu's dataset[7] (GEO accession number: GSE89497). In the original studies, cells have been categorized by their stages and lineages into 7 coarse clusters (3 in pre- to post- implantation, 3 in first-trimester placenta, and 1 in second-trimester placenta; N.B., Zhou's dataset annotation was obtained from Castel. et al.[8]). Here, we performed pseudotime analysis on these two datasets jointly. First, we pre-processed the combined dataset by 1) log-transformation of the TPM counts, 2) scaling to 10 on each gene, and 3) regressing out on the sequencing depth. Then, we calculated the diffusion pseudotime by DPT using scanpy.tl.dpt with default parameters[9]. Cells with inconsistent stage-lineage annotations and pseudotime values have been removed, and 4041 and 952 cells were retained for downstream analysis, respectively. Based on log counts, Pearson correlations were computed for selected markers with ACE2, and visualized by scatter plots. Batch effects among bulk RNA-seq have also been regressed out before comparing in vitro cells to in vivo reference.

**Integration of in vitro and in vivo extra-embryonic datasets**

For integrative analysis, we mapped the in vitro bulk RNA-seq to the in vivo scRNA reference by using singular value decomposition (SVD) modeling[10,11], based on the assumption that top components could capture cell identity and biological variations, regardless of sequencing types. First, the batch effects between the two scRNA datasets were regressed out to construct an extra-embryonic landscape containing peri-implantation to placenta stages. Then, we fitted the SVD model using the whole transcriptome of the combined in vivo scRNA reference, and generated a 50-component decomposition result for the in vivo reference. Next, we applied the fitted model to project the in vitro cells (bulk RNA-seq) to the corresponding 50 components in the same space. After concatenating the bulk RNA and scRNA datasets as 50-component samples, UMAP visualization could be generated. As a proof of effectivity of the SVD modelling in segregating cell types, for the in vivo sector, inter-cell type variations were captured by each cluster in the UMAP.

**Whole-transcriptome correlation analysis between in vitro cell line bulk RNA-seq and in vivo scRNA-seq**

With the assumption that bulk RNA-seq reflect additive effects of scRNA levels, we generated pseudo-bulk sets from the in vivo scRNA-seq. For each in vivo subtype, we randomly selected two sets of 50 cells as two "pseudo-bulk cells" and calculated an average to simulate two bulk samples for each subtype, hence we have 14 pseudo-bulk samples. After merging the 14 pseudo-bulk samples with in vitro cells (19 samples, 10 types), we calculated the pair-wise Pearson coefficients across the whole transcriptome between each two samples. The coefficients were visualized in the cluster map to impute the corresponding in vivo stage of each in vitro cell line.

**Integrative pseudotime analysis**

First, diffusion pseudotime was computed for the whole in vivo reference using SCANPY (scanpy.tl.dpt, default parameters). We then extracted the PI_TB to PI_STB subsets as the pseudotime reference set for downstream analysis of in vitro ST differentiation cells. There are 2295 PI_TB cells and 1282 PI_STB cells, constituting a unidirectional differentiation trajectory. Similar to the process described in Integration of bulk RNA and scRNA datasets part, we fitted a 50-component SVD model using the scRNA reference to reduce the dimensions of both scRNA and bulk RNA to a common 50-component space. We hypothesized that these 50 components could predict the pseudotime calculated by SCANPY. The machine learning prediction process was achieved using sklearn.linear_model.LinearRegression. Briefly, the 50 components and the pseudotime of 1721 PI_TB cells and 961 PI_STB cells (75% of each subtype) were used to train the linear regression model, and the remaining 25% test set of each subtype were used to validate the model. The validation result was visualized by the density plot. After confirming the effectivity of the model, we applied the model to predict the pseudotime of in vitro differentiation cells. We also calculated the theoretical "observed pseudotime" for in vitro cells by the following linear equation between the predicted and observed pseudotime in the in vivo reference:

$$Observed\ pseudotime\ = 1.022 * Predicted\ pseudotime - 0.0032$$

Then pseudotime of each dataset was then merged to visualize the relative differentiation stage of each cell.

#### QUANTIFICATION AND STATISTICAL ANALYSIS

Statistical analysis was carried out with Prism software using one or two tailed unpaired Student t test for comparison of two groups and ANOVA for comparisons of multiple groups. The threshold for statistical significance was $p < 0.05$. All details on sample size, the number of replicates, statistical tests and p values for each experiment are provided in the relevant figures and legends. Unless differently specified in the figure legend, n refers to the number of replicates.
