## Supplemental Tables for "Human Early Syncytiotrophoblasts Are Highly Susceptible to SARS-CoV-2 Infection"

### KEY RESOURCES TABLE

| REAGENT or RESOUCE | SOURCE | IDENTIFIER |
|---|---|---|
| **Antibodies** | | |
| Anti-Oct-3/4 | R and D Systems | AF1759-SP |
| Anti-TEAD4 | Abcam | ab58310 |
| Anti-CDX2 antibody [EPR2764Y] | Abcam | ab76541 |
| GCM1 Antibody | Novus biologicals | NBP2-48520-25ul |
| PE anti-human CD49e Antibody | Biolegend | 328009 |
| Purified anti-human/mouse CD49f Antibody | Biolegend | 313602 |
| FITC anti-human CD49a Antibody | Biolegend | 328307 |
| Alexa Fluor® 488 anti-human HLA-A,B,C Antibody | Biolegend | 311413 |
| ENDOU Antibody | Novus biologicals | NBP2-55877 |
| Anti-E Cadherin [EP700Y] | Abcam | ab40772 |
| Anti-Transcription factor AP-2-alpha [EPR2688(2)] | Abcam | Abcam |
| Purified anti-human CD49e Antibody | Biolegend | 328002 |
| Anti-GATA-3 | R and D Systems | MAB6330 |
| Anti-TRA-1-60 | STEMCELL TECHNOLOGIES | 60064 |
| Anti-ACE2 | Thermo Fisher Scientific | SN0754 |
| ACE2 Rabbit mAb | ABclonal | A4612 |
| Anti-SSEA-4 | STEMCELL TECHNOLOGIES | 60062 |
| Anti-hCGB | Thermo Fisher Scientific | 14-6508-82 |
| Anti-KRT7 | Santa Cruz Biotechnology | 5F282 |
| Anti-AP-2 gamma | R and D Systems | AF5059-SP |
| Anti-HLA-G | Abcam | ab7759 |
| Anti-SDC1 | Abcam | ab181789 |
| Anti-CD46 | BioLegend | 352403 |
| Anti-ZO-1 | Abclonal Technologies | A0659 |
| Anti-KRT18 | R and D Systems | MAB7619 |
| Alexa Fluor® 488 anti-human/mouse SSEA-3 Antibody | Biolegend | Cat. 330305 |
| Alexa Fluor® 488 anti-human/mouse TRA-1-60 Antibody | Biolegend | Cat. 330613 |
| Anti-p63/TP73L | R and D Systems | AF1916 |
| Donkey anti-Mouse 555 | Thermo Fisher Scientific | A-31570 |
| Donkey anti-Mouse 488 | Thermo Fisher Scientific | A-21202 |
| Donkey anti-Goat 555 | Thermo Fisher Scientific | A-21432 |
| Donkey anti-Goat 488 | Thermo Fisher Scientific | A-11055 |
| Donkey anti-Rabbit 594 | Thermo Fisher Scientific | A-21207 |
| Donkey anti-Rabbit 488 | Thermo Fisher Scientific | A-21206 |
| Goat anti-Guinea Pig 647 | Thermo Fisher Scientific | A-21450 |
| Rabbit-anti-SARS-CoV-2-NP serum | In house | Yuan et al, Nature, 2021, 10.1038/s41586-021-03431-4 |
| **Bacterial and virus strains** | | |
| SARS-CoV-2 (HKU-001a) | In house | GenBank accession number: MT230904 |
| MERS-CoV (EMC/2012) | Erasmus Medical Center, Netherlands | GenBank accession no. JX869059.2 |
| **Chemicals, peptide, and recombinant proteins** | | |
| GC376 | MedChemExpress | HY-100721 |
| CTS™ (Cell Therapy Systems) N-2 Supplement | Thermo Fisher | A1370701 |
| B-27™ Supplement (50X), serum free | Thermo Fisher | 17504044 |
| XAV939 | MedChemExpress | HY-15147 |
| Neurobasal Medium | Thermo Fisher | 21103049 |
| Remdesivir | MedChemExpress | HY-104077 |
| DMEM/F-12, no glutamine | Thermo Fisher | 21331020 |
| Y-27632 dihydrochloride | Tocris | 1254/10 |
| FBS | Thermo Fisher | 10270 |
| LIF Recombinant Human Protein | Thermo Fisher | PHC9484 |
| DPBS, powder, no calcium, no magnesium | Thermo Fisher | 21600010 |
| FastQuant RT Super Mix | TIANGEN | KR108-01 |
| PowerUp™ SYBR™ Green Master Mix | Thermo Fisher | A25776 |
| SB-431542 | Tocris | 1614 |
| A83-01 | Tocris | 2939 |
| CHIR-99021 | Tocris | 4423 |
| A 419259 trihydrochloride | MedChemExpress | HY-15764A |
| Geltrex™ LDEV-Free Reduced Growth Factor Basement Membrane Matrix | Thermo Fisher | A1413201 |

| | | |
|---|---|---|
| MicroAmp™ Fast Optical 96-Well Reaction Plate with Barcode, 0.1 mL | Thermo Fisher | 4346906 |
| MicroAmp™ Optical Adhesive Film | Thermo Fisher | 4360954 |
| Forskolin | Sigma | F3917 |
| Matrigel Matrix. GFR | Corning | 354230 |
| EGF Recombinant Human Protein | Thermo Fisher | PHG0311 |
| Prostaglandin E2 | MedChemExpress | HY-101952 |
| HGF Protein, Human, Recombinant | Sino biological | Cat: 10463-HNAS |
| RSPO1 Protein, Human, Recombinant | Sino biological | Cat: 11083-HNAS |
| DMSO, Anhydrous | Thermo Fisher | D12345 |
| Bovine Serum Albumin solution | Thermo Fisher | 15260037 |
| Recombinant Human NRG1-beta 1/HRG1-beta 1 EGF Domain Protein | R&D | 396-HB-050 |
| Normal Donkey Serum | Abcam | ab7475 |
| KO serum replacement | Thermo Fisher | 10828010 |
| Valproic Acid (Sodium Salt) | Stem cell technology | 72292 |
| Cell Recovery Solution | Corning | 354253 |
| Recombinant Human FGF basic/FGF2/bFGF (146 aa) Protein | R&D | 233-FB-500/CF |
| µ-Slide 8 Well high Glass Bottom | ibidi | 80807 |
| b-Mercaptoethanol | Thermo Fisher | 31350010 |
| TryPLE-Express | Thermo Fisher | 12605036 |
| Trypsin/EDTA | Thermo Fisher | 25200072 |
| Penicillin-Streptomycin-Glutamine (100X) | Thermo Fisher | 10378016 |
| Minimum Essential Medium (MEM) Vitamin Solution | Thermo Fisher | 11120052 |
| Insulin-Transferrin-Selenium-Ethanolamine (ITS-X) (100X) | Thermo Fisher | 51500056 |
| Triton X-100 | Sigma | T9284 |
| 4% Paraformaldehyde | Sigma | P6148 |
| L-Ascorbic acid | Sigma | 49752-100G |
| **Critical commercial assays** | | |
| CellTiter-Glo® Luminescent Cell Viability Assay | Promega | G7570 |
| miRcute Plus miRNA cDNA First-Strand cDNA Kit | TIANGEN | KR211-01 |
| miRcute Plus miRNA qPCR Kit (SYBR Green) | TIANGEN | FP411-01 |
| QuantiNova Probe RT-PCR kit | QIAGEN | 208354 |
| Transwell invasion assay | Corning | 354480 |
| **Deposited data** | | |
| RNA-seq data | (Okae, H.,2018) | JGA: JGA00000000074<br>JGA: JGA00000000117<br>JGA: JGA00000000122 |
| RNA-seq data | (Gao, X.,2019) | E-MTAB-7253 |
| RNA-seq data | (Sheridan, 2021) | E-MTAB-10429 |
| scRNA-seq data | (Zhou, F. et al., 2019) | GSE109555 |
| scRNA-seq data | (Liu, Y. et al., 2018) | GSE89497 |
| RNA-seq data | This paper | GSE190432 |
| **Experimental models: cell lines** | | |
| Monkey: Vero E6 cells | ATCC | CCL-81 |
| Human embryonic stem cell (hESC) line: Man-1/M1 | (Camarasa, 2010) | N/A |
| Caco-2 [Caco2] | ATCC | HTB-37 |
| TSC-BST, Human Blastocyst derived hTSCs | (Okae et al., 2018) | N/A |
| **Oligonucleotides** | | |
| See Methods Table S3 for the sequences of oligonucleotides used in this study | This paper | Methods Table S3 |
| **Software and algorithms** | | |
| ImageJ Fiji (2.0.0) | NIH | https://imagej.net/Fiji |
| GraphPad Prism 8.0 | GraphPad | https://www.graphpad.com/scientific-software/prism/ |

| | | |
|---|---|---|
| Microsoft | Microsoft | https://www.microsoft.com/de-at/microsoft-365/excel |
| Adobe Illustrator | Adobe | https://www.adobe.com/at/products/illustrator.html |
| FlowJo | BD Life Sciences | https://www.flowjo.com/ |
| Zeiss Zen (Blue edition) | Zeiss | https://www.zeiss.com/microscopy/int/products/microscope-software/zen-lite.html |
| FeatureCounts v2.0.1 | N/A | http://subread.sourceforge.net |
| SCANPY (v1.7.2, scRNA /integrative analysis) | N/A | https://scanpy-tutorials.readthedocs.io/en/latest/index.html |

### SUPPLEMENTAL ITEMS

#### Table S1. Amnion and trophoblast signature genes[12]. Related to Figure1.

| gene | Type | annotaton |
|---|---|---|
| CA3 | Amnion | carbonic anhydrase 3 |
| DIRAS2 | Amnion | DIRAS family GTPase 2 |
| FXYD6 | Amnion | FXYD domain containing ion transport regulator 6 |
| GABRP | Amnion | gamma-aminobutyric acid type A receptor pi subunit |
| HEPH | Amnion | hephaestin |
| IGFBP5 | Amnion | insulin like growth factor binding protein 5 |
| ISL1 | Amnion | ISL LIM homeobox 1 |
| LMO1 | Amnion | LIM domain only 1 |
| LRRN1 | Amnion | leucine rich repeat neuronal 1 |
| MSRB2 | Amnion | methionine sulfoxide reductase B2 |
| RARRES2 | Amnion | retinoic acid receptor responder 2 |
| SDC2 | Amnion | syndecan 2 |
| CDX2 | Amnion | caudal type homeobox 2 |
| MUC16 | Amnion | mucin 16, cell surface associated |
| ITGB6 | Amnion | integrin subunit beta 6 |
| HAND1 | Amnion | heart and neural crest derivatives expressed 1 |
| SEMA3C | Amnion | semaphorin 3C |
| PMP22 | Amnion | peripheral myelin protein 22 |
| TRIM55 | Amnion | tripartite motif containing 55 |
| AC011453.1 | TE | NA |
| ADAM15 | TE | ADAM metallopeptidase domain 15 |
| AEN | TE | apoptosis enhancing nuclease |
| AGPAT5 | TE | 1-acylglycerol-3-phosphate O-acyltransferase 5 |
| AKAP13 | TE | A-kinase anchoring protein 13 |
| AKAP8 | TE | A-kinase anchoring protein 8 |
| ATP13A3 | TE | ATPase 13A3 |
| ATXN2L | TE | ataxin 2 like |
| B4GALT1 | TE | beta-1,4-galactosyltransferase 1 |
| BAIAP2 | TE | BAI1 associated protein 2 |
| BOP1 | TE | block of proliferation 1 |
| BUB1 | TE | BUB1 mitotic checkpoint serine/threonine kinase |
| BYSL | TE | bystin like |
| C6orf106 | TE | chromosome 6 open reading frame 106 |
| CARS | TE | cysteinyl-tRNA synthetase |
| CCDC86 | TE | coiled-coil domain containing 86 |
| CCKBR | TE | cholecystokinin B receptor |
| CCNA2 | TE | cyclin A2 |
| CCND3 | TE | cyclin D3 |
| CCR7 | TE | C-C motif chemokine receptor 7 |
| CD3EAP | TE | CD3e molecule associated protein |
| CDC6 | TE | cell division cycle 6 |
| CDCA4 | TE | cell division cycle associated 4 |
| CDK12 | TE | cyclin dependent kinase 12 |
| CEP85 | TE | centrosomal protein 85 |
| CHERP | TE | calcium homeostasis endoplasmic reticulum protein |
| CYP11A1 | TE | cytochrome P450 family 11 subfamily A member 1 |
| CYP19A1 | TE | cytochrome P450 family 19 subfamily A member 1 |
| DCAF12 | TE | DDB1 and CUL4 associated factor 12 |
| DDX54 | TE | DEAD-box helicase 54 |
| DEPP1 | TE | NA |
| DHX38 | TE | DEAH-box helicase 38 |
| DLG5 | TE | discs large MAGUK scaffold protein 5 |
| DLX3 | TE | distal-less homeobox 3 |
| DNAJA3 | TE | DnaJ heat shock protein family (Hsp40) member A3 |
| DNM2 | TE | dynamin 2 |

| | | |
|---|---|---|
| DNMT3B | TE | DNA methyltransferase 3 beta |
| DNMT3L | TE | DNA methyltransferase 3 like |
| DPH2 | TE | DPH2 homolog |
| DPPA3 | TE | developmental pluripotency associated 3 |
| EAF1 | TE | ELL associated factor 1 |
| ECPAS | TE | NA |
| EHD1 | TE | EH domain containing 1 |
| EHD4 | TE | EH domain containing 4 |
| ELMSAN1 | TE | ELM2 and Myb/SANT domain containing 1 |
| EP300 | TE | E1A binding protein p300 |
| ERVW-1 | TE | endogenous retrovirus group W member 1, envelope |
| FAM98A | TE | family with sequence similarity 98 member A |
| FSCN1 | TE | fascin actin-bundling protein 1 |
| FURIN | TE | furin, paired basic amino acid cleaving enzyme |
| FYB1 | TE | NA |
| GABARAPL1 | TE | GABA type A receptor associated protein like 1 |
| GATA2 | TE | GATA binding protein 2 |
| GATAD2A | TE | GATA zinc finger domain containing 2A |
| GCM1 | TE | glial cells missing homolog 1 |
| GCN1 | TE | GCN1, eIF2 alpha kinase activator homolog |
| GDE1 | TE | glycerophosphodiester phosphodiesterase 1 |
| GEMIN4 | TE | gem nuclear organelle associated protein 4 |
| GJA5 | TE | gap junction protein alpha 5 |
| GNPNAT1 | TE | glucosamine-phosphate N-acetyltransferase 1 |
| GREM2 | TE | gremlin 2, DAN family BMP antagonist |
| GSE1 | TE | Gse1 coiled-coil protein |
| GTSF1 | TE | gametocyte specific factor 1 |
| GYS1 | TE | glycogen synthase 1 |
| HCFC1 | TE | host cell factor C1 |
| HK2 | TE | hexokinase 2 |
| HMOX1 | TE | heme oxygenase 1 |
| KNL1 | TE | kinetochore scaffold 1 |
| KNSTRN | TE | kinetochore localized astrin/SPAG5 binding protein |
| LASP1 | TE | LIM and SH3 protein 1 |
| LCMT1-AS2 | TE | NA |
| LETM1 | TE | leucine zipper and EF-hand containing transmembrane protein 1 |
| LINC00668 | TE | NA |
| LMNB2 | TE | lamin B2 |
| LRRC58 | TE | leucine rich repeat containing 58 |
| LYAR | TE | Ly1 antibody reactive |
| MAK16 | TE | MAK16 homolog |
| MBD3 | TE | methyl-CpG binding domain protein 3 |
| MCM10 | TE | minichromosome maintenance 10 replication initiation factor |
| MED12L | TE | mediator complex subunit 12 like |
| MED13 | TE | mediator complex subunit 13 |
| METTL13 | TE | methyltransferase like 13 |
| MFN2 | TE | mitofusin 2 |
| MIEF1 | TE | mitochondrial elongation factor 1 |
| MKI67 | TE | marker of proliferation Ki-67 |
| MRGPRX1 | TE | MAS related GPR family member X1 |
| MTHFD2 | TE | methylenetetrahydrofolate dehydrogenase (NADP+ dependent) 2, |
| MTIF2 | TE | mitochondrial translational initiation factor 2 |
| MYBBP1A | TE | MYB binding protein 1a |
| NABP1 | TE | nucleic acid binding protein 1 |
| NAT10 | TE | N-acetyltransferase 10 |
| NCLN | TE | nicalin |
| NCOA3 | TE | nuclear receptor coactivator 3 |
| NID1 | TE | nidogen 1 |
| NLRP2 | TE | NLR family pyrin domain containing 2 |
| NLRP7 | TE | NLR family pyrin domain containing 7 |
| NOL10 | TE | nucleolar protein 10 |

| | | |
|---|---|---|
| NOL6 | TE | nucleolar protein 6 |
| NOP2 | TE | NOP2 nucleolar protein |
| NUP98 | TE | nucleoporin 98 |
| OGT | TE | O-linked N-acetylglucosamine (GlcNAc) transferase |
| OVOL1 | TE | ovo like transcriptional repressor 1 |
| PFAS | TE | phosphoribosylformylglycinamidine synthase |
| PGF | TE | placental growth factor |
| PIM1 | TE | Pim-1 proto-oncogene, serine/threonine kinase |
| PIP5K1A | TE | phosphatidylinositol-4-phosphate 5-kinase type 1 alpha |
| PKMYT1 | TE | protein kinase, membrane associated tyrosine/threonine 1 |
| PMM2 | TE | phosphomannomutase 2 |
| PNP | TE | purine nucleoside phosphorylase |
| POLR3E | TE | RNA polymerase III subunit E |
| PPIF | TE | peptidylprolyl isomerase F |
| PPP1R10 | TE | protein phosphatase 1 regulatory subunit 10 |
| PPRC1 | TE | peroxisome proliferator-activated receptor gamma, coactivator-related 1 |
| PRPS2 | TE | phosphoribosyl pyrophosphate synthetase 2 |
| PRRC2B | TE | proline rich coiled-coil 2B |
| PSME4 | TE | proteasome activator subunit 4 |
| PTDSS1 | TE | phosphatidylserine synthase 1 |
| PTGES | TE | prostaglandin E synthase |
| RAB11FIP4 | TE | RAB11 family interacting protein 4 |
| RAB35 | TE | RAB35, member RAS oncogene family |
| RACGAP1 | TE | Rac GTPase activating protein 1 |
| RANGAP1 | TE | Ran GTPase activating protein 1 |
| RBM14 | TE | RNA binding motif protein 14 |
| RBM47 | TE | RNA binding motif protein 47 |
| RCC1 | TE | regulator of chromosome condensation 1 |
| RCL1 | TE | RNA terminal phosphate cyclase like 1 |
| RDH13 | TE | retinol dehydrogenase 13 |
| REEP1 | TE | receptor accessory protein 1 |
| RHOBTB1 | TE | Rho related BTB domain containing 1 |
| RHOG | TE | ras homolog family member G |
| RHOT2 | TE | ras homolog family member T2 |
| RNF168 | TE | ring finger protein 168 |
| RP2 | TE | RP2, ARL3 GTPase activating protein |
| RRM2 | TE | ribonucleotide reductase regulatory subunit M2 |
| RRP12 | TE | ribosomal RNA processing 12 homolog |
| RRS1 | TE | ribosome biogenesis regulator homolog |
| RYBP | TE | RING1 and YY1 binding protein |
| S1PR2 | TE | sphingosine-1-phosphate receptor 2 |
| SDC1 | TE | syndecan 1 |
| SENP5 | TE | SUMO1/sentrin specific peptidase 5 |
| SESN2 | TE | sestrin 2 |
| SLC1A3 | TE | solute carrier family 1 member 3 |
| SLC1A5 | TE | solute carrier family 1 member 5 |
| SLC35F6 | TE | solute carrier family 35 member F6 |
| SLC40A1 | TE | solute carrier family 40 member 1 |
| SLC4A2 | TE | solute carrier family 4 member 2 |
| SLC7A5 | TE | solute carrier family 7 member 5 |
| SLC7A6 | TE | solute carrier family 7 member 6 |
| SMG5 | TE | SMG5, nonsense mediated mRNA decay factor |
| SNX8 | TE | sorting nexin 8 |
| SOAT1 | TE | sterol O-acyltransferase 1 |
| SP6 | TE | Sp6 transcription factor |
| SRCAP | TE | Snf2 related CREBBP activator protein |
| SRRT | TE | serrate, RNA effector molecule |
| ST6GAL1 | TE | ST6 beta-galactoside alpha-2,6-sialyltransferase 1 |
| STS | TE | steroid sulfatase |
| SURF6 | TE | surfeit 6 |
| TEAD1 | TE | TEA domain transcription factor 1 |

| | | |
|---|---|---|
| TFRC | TE | transferrin receptor |
| TGFBR3 | TE | transforming growth factor beta receptor 3 |
| TIGAR | TE | TP53 induced glycolysis regulatory phosphatase |
| TLE3 | TE | transducin like enhancer of split 3 |
| TMEM109 | TE | transmembrane protein 109 |
| TOB2 | TE | transducer of ERBB2, 2 |
| TRAF4 | TE | TNF receptor associated factor 4 |
| TRIP12 | TE | thyroid hormone receptor interactor 12 |
| TRIP13 | TE | thyroid hormone receptor interactor 13 |
| TYRO3 | TE | TYRO3 protein tyrosine kinase |
| UBR4 | TE | ubiquitin protein ligase E3 component n-recognin 4 |
| USP5 | TE | ubiquitin specific peptidase 5 |
| UTP20 | TE | UTP20, small subunit processome component |
| VARS | TE | valyl-tRNA synthetase |
| ZFHX3 | TE | zinc finger homeobox 3 |

### Table S2. Correlation index between ACE2 and Trophoblast markers，related to Figure2.

| | coefficient | p_value |
|---|---|---|
| CDX2 | 0.04266543 | 0.00256635 |
| CD46 | 0.24451753 | 7.03E-69 |
| TFRC | -0.0493468 | 0.00048636 |
| GCM1 | -0.0636706 | 6.72E-06 |
| CGB3 | 0.10253989 | 3.79E-13 |
| CGB5 | 0.19950932 | 5.37E-46 |
| SDC1 | 0.10222731 | 4.46E-13 |
| GATA2 | -0.0356747 | 0.01170267 |
| GATA3 | -0.0968373 | 7.03E-12 |
| TP63 | -0.0269452 | 0.05692952 |
| TEAD4 | -0.1328758 | 4.15E-21 |
| ACE2 | 1 | 0 |
| ENG | 0.20688926 | 2.11E-49 |
| ITGA5 | 0.10266201 | 3.55E-13 |
| ITGA6 | -0.0676585 | 1.71E-06 |
| CSH1 | 0.18957838 | 1.27E-41 |
| CSH2 | 0.24808141 | 6.54E-71 |
| HLA-G | 0.0706351 | 5.85E-07 |
| MMP2 | 0.03941273 | 0.00534726 |

### Table S3. List of PCR primers

| Gene Name | Forward (5’-3’) | Reverse (5’-3’) |
|---|---|---|
| SARS-CoV-2 | CGCATACAGTCTTRCAGGCT | GTGTGATGTTGAWATGACATGGTC |
| ACE2 (Exon17-18) | GGAGTTGTGATGGGAGTGAT | GATGGAGGCATAAGGATTTT |
| ACE2 (Exon9-10) | TCCATTGGTCTTCTGTCACCCG | AGACCATCCACCTCCACTTCTC |
| TMPRSS2 | CTCTACGGACCAAACTTCATC | CCACTATTCCTTGGCTAGAGTA |
| CD147 | GGCTGTGAAGTCGTCAGAACAC | ACCTGCTCTCGGAGCCGTTCA |
| NANOG | TGAACCTCAGCTACAAACAG | TGGTGGTAGGAAGAGTAAAG |
| OCT4 | CCTCACTTCACTGCACTGTA | CAGGTTTTCTTTCCCTAGCT |
| SOX2 | TTCACATGTCCCAGCACTACCAGA | TCACATGTGTGAGAGGGGCAGTGTGC |
| CDX2 | TTCACTACAGTCGCTACATCACC | TTGATTTTCCTCTCCTTTGCTC |
| GATA3 | ACATCTCGCCCTTCAGCCAC | CATGGCGGTGACCATGCTGGA |
| KRT7 | AGGATGTGGATGCTGCCTAC | CACCACAGATGTGTCGGAGA |
| TEAD4 | CAGGTGGTGGAGAAAGTTGAGA | GTGCTTGAGCTTGTGGATGAAG |
| TFAP2C | ACAGGATCCATGTTGTGGAAAATAACCGAT | ATACTCGAGTTTCCTGTGTTTCTCCATTTT |
| TP63 | AGAAACGAAGATCCCCAGATGA | CTGTTGCTGTTGCCTGTACGTT |
| CGB | ACCCTGGCTGTGGAGAAGG | ATGGACTCGAAGCGCACA |
| ERVW-1 | GTTAATGACATCAAAGGCACCC | CCCCATCTCAACAGGAAAACC |
| SDC1 | GCTGACCTTCACACTCCCCA | CAAAGGTGAAGTCCTGCTCCC |
| HLA-G | CAGATACCTGGAGAACGGGA | CAGTATGATCTCCGCAGGGT |
| MMP2 | TGGCACCCATTTACACCTACAC | ATGTCAGGAGAGGCCCCATAGA |
| ITGB6 | CTCAACACAATAAAGGAGCTGGG | AAAGGGGATACAGGTTTTTCCAC |
| GABRP | TTTCTCAGGCCCAATTTTGGT | GCTGTCGGAGGTATATGGTGG |
| MUC16 | GGAGCACACGCTAGTTCAGAA | GGTCTCTATTGAGGGGAAGGT |
| VTCN1 | TCTGGGCATCCCAAGTTGAC | TCCGCCTTTTGATCTCCGATT |
| cGAS | TAACCCTGGCTTTGGAATCAAAA | TGGGTACAAGGTAAAATGGCTTT |
| ZBP1 | TGGTCATCGCCCAAGCACTG | GGCGGTAAATCGTCCATGCT |

| | | |
|---|---|---|
| MDA5 | GAGCAACTTCTTTCAACCACAG | CACTTCCTTCTGCCAAACTTG |
| STING1 | AGCATTACAACAACCTGCTACG | GTTGGGGTCAGCCATACTCAG |
| IFNA2 | CTTGAAGGACAGACATGACTTTGGA | GGATGGTTTCAGCCTTTTGGA |
| IFNB1 | AAACTCATGAGCAGTCTGCA | AGGAGATCTTCAGTTTCGGAGG |
| IFNG | TGGCTTTTCAGCTCTGCATC | CCGCTACATCTGAATGACCTG |
| IFNL1 | CGCCTTGGAAGAGTCACTCA | GAAGCCTCAGGTCCCAATTC |
| IFNL2 | AGTTCCGGGCCTGTATCCAG | GAGCCGGTACAGCCAATGGT |
| IFNL3 | TCGCTTCTGCTGAAGGACTGCA | CCTCCAGAACCTTCAGCGTCAG |
| IFNL4 | ATGCGGCCGAGTGTCTGG | GCTCCAGCGAGCGGTAGTG |
| IL6 | GTCAGGGGTGGTTATTGCAT | AGTGAGGAACAAGCCAGAGC |
| IL28A | TCCAGTCACGGTCAGCA | CAGCCTCAGAGTGTTTCTTCT |
| TNF | CTCTTCTGCCTGCTGCACTTTG | ATGGGCTACAGGCTTGTCACTC |
| HLA-A | CGAGGATGGCCGTCATGGCG | CACATTCCGTGTCTCCTGGTCCC |
| HLA-B | CAGTTCGTGAGGTTCGACAG | CAGCCGTACATGCTCTGGA |
| ITGA1 | CTGGACATAGTCATAGTGCTGGA | ACCTGTGTCTGTTTAGGACCA |
| ITGA5 | GTCGGGGGCTTCAACTTAGAC | CCTGGCTGGCTGGTATTAGC |
| ITGA6 | CACATCTCCTCCCTGAGCAC | TATCTTGCCACCCATCCTTG |
| CDX2 | TTCACTACAGTCGCTACATCACC | TTGATTTTCCTCTCCTTTGCTC |
| ELF5 | TGCCCTCACGGTAATGTTGGA | TGATGCTCAAAGGCAGGGTAG |
| GAPDH | CAAATTCCATGGCACCGTCA | ATCGCCCCACTTGATTTTGG |
| human miR-103a | GTAGCAGCATTGTACAGGG | |
| human miR-526b-3p | GTTTGGGAAAGTGCTTCCTTTT | |
| human miR-517a | GTTTGGATCGTGCATCCTTTTA | |
| human miR-517b | GTGCCTCTAGATGGAAGCA | |
| human miR-525-3p | GTTGAAGGCGCTTCCCTTT | |
